# Activating natural product synthesis using CRISPR interference and activation systems in *Streptomyces*

**DOI:** 10.1101/2021.10.28.466254

**Authors:** Andrea Ameruoso, Maria Claudia Villegas Kcam, Katherine Piper Cohen, James Chappell

## Abstract

The rise of antibiotic-resistant bacteria represents a major threat to global health, creating an urgent need to discover new antibiotics. Natural products derived from the genus *Streptomyces* represent a rich and diverse repertoire of chemical molecules from which new antibiotics are likely to be found. However, a major challenge is that the biosynthetic gene clusters (BGCs) responsible for natural product synthesis are often poorly expressed under laboratory culturing conditions, thus preventing isolation and screening of novel chemicals. To address this, we describe a novel approach to activate silent BGCs through rewiring endogenous regulation using synthetic gene regulators based upon CRISPR-Cas. First, we create CRISPR interference (CRISPRi) and CRISPR activation (CRISPRa) systems that allow for highly programmable and effective gene repression and activation in *Streptomyces*. We then harness these tools to activate a silent BGC through perturbing its endogenous regulatory network. Together, this work advances the synthetic regulatory toolbox for *Streptomyces* and facilitates the programmable activation of silent BGCs for novel chemical discovery.

## INTRODUCTION

Bacterial resistance to antimicrobial agents is globally recognized as one of the major challenges facing public health^1,2^. Addressing the antibiotic resistance crisis requires a multipronged approach, including the discovery of new antibiotics for which resistance has not been reported^3^. However, over the last decades, there has been a sharp decline in the discovery of new antibiotics, and it is widely recognized that innovative approaches for antibiotic discovery are required^4^.

Natural products derived from *Streptomyces* species are a likely major source of new antibiotics^5^. Microbial natural products are encoded by convoluted genomic regions called Biosynthetic Gene Clusters (BGCs). BGCs typically consist of dozens of genes that are co-localized at a single portion of a bacterial genome and encode components for natural product synthesis, transport, regulation, and resistance^6^. While natural products derived from *Streptomyces* species were once thought to be exhausted, advances in bioinformatic tools and genome sequencing have revealed that BGCs are far more abundant than previously thought, averaging 39 per genome^6^. Additionally, greater diversity is likely to exist: evidence indicates that the same species can vary tremendously in the BGCs they carry, due to high-rates of horizontal gene transfer and formation of novel clusters through duplication and rearrangement events^6^. Furthermore, new natural products are likely to come from exploring *Streptomyces* species beyond traditionally mined soil microbiomes. For example, *Streptomyces* have been found within host-associated microbiomes of higher-order eukaryotes such as humans^7^, marine organisms^8^ and insects^9^, which are likely to contain different chemical diversity due to distinct evolutionary trajectories and environmental ecologies^10^.

While a rich repertoire of BGCs exist, the vast majority remain uncharacterized^11^. A key reason for this lack of extensive characterization is that the majority of BGCs are expressed poorly or not at all when grown under laboratory conditions^12^. This is due to stringent expression control imposed by networks of gene regulators located within the BGC, called cluster-situated regulators (CSRs), and/or global regulators encoded at distinct genomic loci. These networks ensure that BGCs are only expressed under certain environmental stimuli^13^. Identifying these stimuli to induce BGC expression in the laboratory requires screening of large parameter spaces^10^. Additionally, it is hard to activate specific and individual BGCs using this strategy^5^. Taken together, there is strong motivation to find new synthetic routes to predictably and precisely activate silent BGCs.

Engineering of CRISPR-Cas systems has led to a powerful suite of *trans*-acting gene regulatory tools able to precisely program gene expression in bacteria^14–16^. For example, CRISPR interference (CRISPRi) leverages a catalytically inactive version of the *Streptococcus pyogenes* Cas9 (dead Cas9, or dCas9) and a single guide RNA (sgRNA) to target DNA to sterically block transcription initiation and elongation^17,18^. In addition to gene repression, CRISPR activation (CRISPRa) systems able to turn on transcription of a target gene have also been created^18–23^. CRISPR activation is achieved by recruiting protein activation domains (ADs) to the dCas9 complex that stimulate transcription when localized upstream of promoter elements. The power of these regulatory tools lies in the facile synthesis and flexibility of sgRNAs, which can be designed to target any DNA sequence that is proximal to a three-nucleotide protospacer adjacent motif (PAM). Additionally, because these systems act *in trans*, they can be encoded on easily-transferable plasmids and provide a route to perturb the expression of chromosome-encoded genes without the need for chromosome engineering that can be time-consuming and arduous. Taken together, CRISPRi and CRISPRa provide a strategy to rapidly and precisely perturb the expression of endogenous genes.

We posit that CRISPRi and CRISPRa can provide a novel approach to activate BGC expression through perturbing and rewiring the underlying regulatory gene networks. For example, CRISPRa could be used to directly activate BGCs or to induce overexpression of endogenous transcription activators controlling BGC expression. Likewise, CRISPRi could be used to relieve BGC repression exerted by endogenous regulators. Importantly, the CRISPRi regulatory mechanism is highly portable, and CRISPRi tools have been developed for *Streptomyces* species. While these tools have been applied to turn off BGC expression^24–26^ and redirect metabolic flux from primary to secondary metabolism^27^, their application to activate silent BGC has yet to be explored. As for CRISPRa, its development has proven especially challenging in bacteria, and the majority of efforts have focused on optimizing these technologies for the model species *E. coli*^18–23^. Crucially, CRISPRa systems have not been demonstrated in non-model species with high biosynthetic potential, such as *Streptomyces*, or applied to activate natural product synthesis.

To address this, here we applied CRISPRi and CRISPRa to the activation of BGCs in *Streptomyces venezuelae* (*S. venezuelae*). We first characterized the performance of CRISPRi by exploring key parameters with a fluorescent reporter. Next, we established a functional CRISPRa system and explored its design rules. Finally, we demonstrated the applicability of both CRISPR tools for the activation of a silent BGC. Overall, our work introduces a facile and predictable strategy for the activation of silent BGCs within *Streptomyces*.

## RESULTS

### Creating a CRISPRi platform for programmable transcription repression in *Streptomyces venezuelae*

As a starting point, our goal was to establish a CRISPRi system optimized for performance in *S. venezuelae*, which is an increasingly important biotechnological strain rich in silent BGCs^28^. The most established CRISPRi system leverages a dCas9 derived from *S. pyogenes* and a synthetic sgRNA^17^. To achieve repression, the sgRNA can be designed to bind the target gene’s promoter, preventing transcription initiation, and thus turning off the expression of the gene (Figure 1a). Alternatively, this complex can be targeted to the coding sequence of a gene to sterically block transcription elongation (Figure 1b). To initially characterize CRISPRi in *S. venezuelae*, we constructed plasmids that express a codon-optimized dCas9 and a sgRNA from the constitutive promoters rpsL(XC) and gapdh(EL), respectively. To measure transcription repression, we created a reporter strain in which a constitutive mCherry expression cassette, under the control of the KasOp* promoter, was integrated into the ΦC31 site of *S. venezuelae* as previously described^29^. We designed and cloned a corresponding sgRNA to target a PAM located at 11 bp on the non-template strand within the mCherry reporter gene. Additionally, a plasmid containing only the antibiotic resistance cassette was constructed to be used as a no-CRISPRi control. We conjugated these plasmids into the *S. venezuelae* reporter strain and measured mCherry fluorescence (587 nm excitation and 610 nm emission) and optical density at 600 nm for each culture. From these experiments we observed that compared to the no-CRISPRi control, CRISPRi produced significant repression of fluorescence (Figure 1b).

**Figure 1.**
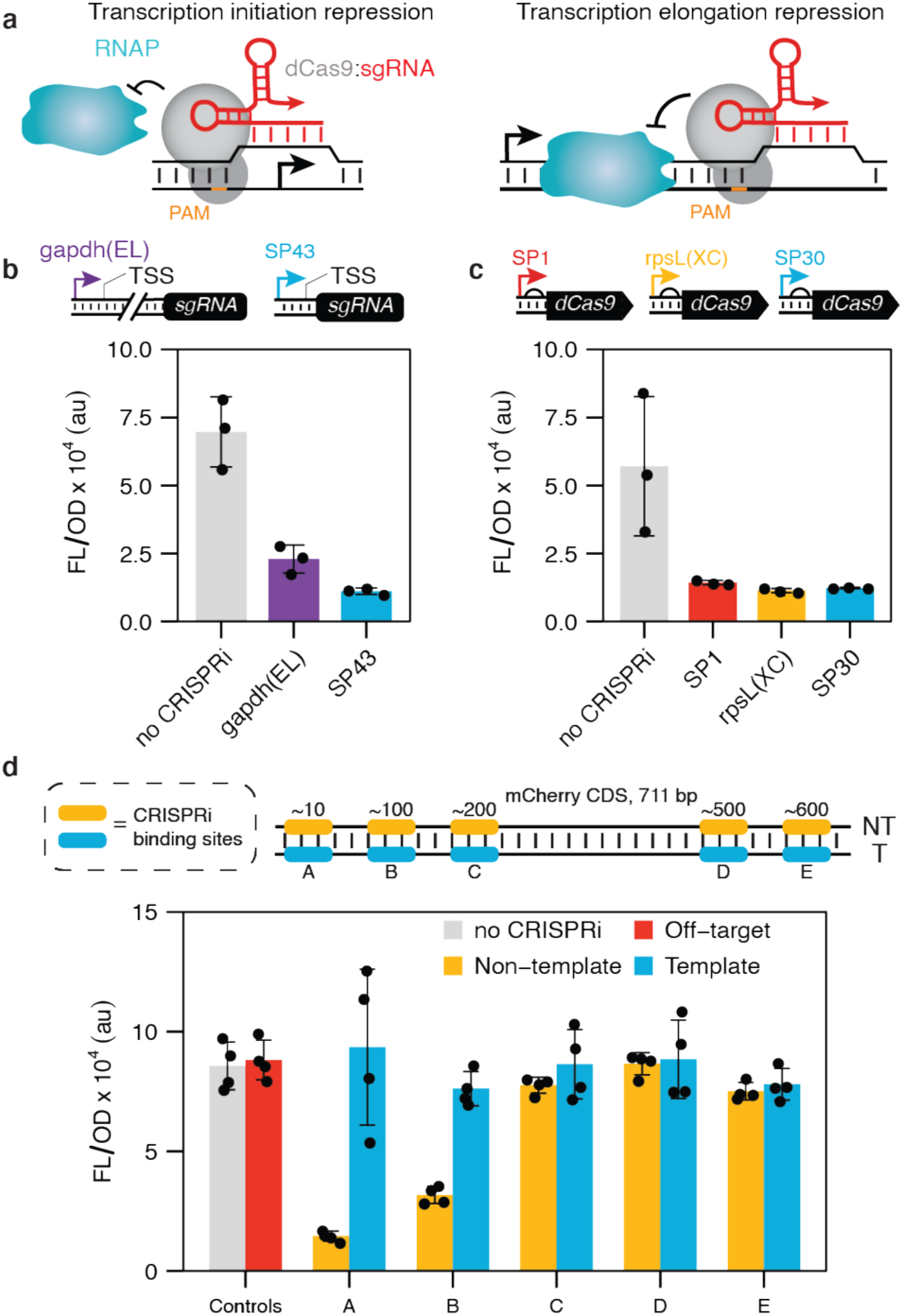
Creating a CRISPR interference (CRISPRi) system for *Streptomyces venezuelae*. **a** Schematic of CRISPRi mechanism. The ribonucleoprotein complex formed by dCas9 (colored *grey*) and the single guide RNA (sgRNA, colored *red*) binds to the promoter or gene coding region to block transcription initiation or elongation by RNA polymerase (RNAP). **b** Eliminating intervening nucleotides between the promoter and the sgRNA leads to higher repression by CRISPRi. Schematic of sgRNA expression cassette with the gapdh(EL) and SP43 promoter. The SP43 promoter contains an annotated TSS, whereas gapdh(EL) TSS remains unannotated. Fluorescence characterization of *S. venezuelae* cells conjugated with CRISPRi plasmid variants designed to repress transcription of a genomically integrated mCherry reporter. **c**. CRISPRi produces robust repression independently of dCas9 expression. Schematic of dCas9 expression cassette using the SP1, rpsL(XC), and SP30 promoters (ascending strength). Fluorescence characterization of *S. venezuelae* cells conjugated with CRISPRi plasmid variants designed to repress transcription of a genomically integrated mCherry reporter. **d**. Position-dependent repression by CRISPRi. Schematic of the sgRNA binding sites used that target different PAMs in the non-template (NT) and template (T) strand of the mCherry reporter gene. Fluorescence characterization of *S. venezuelae* cells conjugated with CRISPRi plasmid variants. Fluorescence characterization was performed by bulk fluorescence measurements (measured in units of fluorescence [FL]/optical density [OD] at 600 nm). Data represent mean values and errors bars represent standard deviation of at least 3 biological replicates.

While successful, we next sought to optimize the performance of CRISPRi by tuning the expression of its components. To do this, we first constructed a library of natural and synthetic constitutive promoters and characterized their expression strength through mCherry fluorescence assays (Supplementary figure 1). Subsequently, we decided to replace the gapdh(EL) promoter driving the expression of the sgRNA with the SP43 promoter^30^. SP43 is not only stronger than gapdh(EL) but has an annotated transcription start site (TSS) that allows for precise expression of the sgRNA without amending additional promoter-encoded sequences onto this transcript. When comparing the two conditions in mCherry fluorescence assays, we observed greater reduction in fluorescence in the presence of SP43 than in the presence of gapdh(EL), indicating stronger CRISPRi repression (Figure 1b). Next, we investigated the effect of dCas9 expression by replacing the original rspL(XC) promoter with a weaker (SP1) and stronger (SP30) promoter (Supplementary figure 1), while keeping the sgRNA under the control of SP43. We observed little difference in the level of transcription repression in response to different dCas9 expression levels (Figure 1c). Interestingly, while performing these experiments we observed a significant decrease in fluorescence in two cases when dCas9 was present: in absence of a sgRNA or in the presence of a non-targeting sgRNA (i.e. a sgRNA designed to target a DNA sequence absent in *Streptomyces*) (Supplementary figure 2). In contrast, inclusion of dCas9 alongside a sgRNA targeting a non-coding region in the genome but not mCherry, resulted in no decrease in fluorescence when compared to a no-CRISPRi control (Supplementary figure 2). Characterizing the growth rate of these control conditions uncovered significant growth defects when dCas9 is present either with a non-targeting sgRNA or without a sgRNA (Supplementary figure 3). While the exact cause of this effect is unknown, we reason they are likely due to nonspecific binding of dCas9 to DNA^31^ that appears to be particularly detrimental for *Streptomyces*, potentially due to the high frequency of PAMs within its GC rich genome^32^.

Having optimized a CRISPRi platform for *S. venezuelae* we next sought to gain deeper insight into the design rules. Specifically, we were interested in understanding how the strand (i.e., non-template or template) and position of CRISPRi targeting within the coding sequence affected the level of transcription repression. To test this, we designed and cloned a series of sgRNAs targeting PAMs in the non-template and template strand of mCherry located at 11, 123, 230, 531, 623, bp and 29, 132, 242, 560, 647 bp respectively. From these experiments we saw significant repression when CRISPRi is targeted to the 5’ end of the coding sequence on the non-template strand and negligible repression when targeting either the template strand or downstream regions of the non-template strand (Figure 1d). This is consistent with previous results of CRISPRi in other microbes^33^.

### Creating a CRISPRa platform for transcription activation in *S. venezuelae*

While CRISPRa systems have been developed for *E. coli*, their application in other bacteria is lacking. For example, to date there has only been a single demonstration in a gram-positive species, *Bacillus subtilis*^34^. To address this, our goal was to establish a CRISPRa platform in *Streptomyces* for the first time. The most established CRISPRa design motif relies upon translationally fusing dCas9 to a protein-based activation domain (AD) that, when localized close to promoter elements, activates transcription through recruitment of the RNAP or stabilization of the RNAP initiation complex (Figure 2a). Variations of this design include use of different ADs, linker sequences, and recruitment strategies^18,19,21,22^. To create a CRISPRa system, we first sought to identify a functional AD for *S. venezuelae*. Specifically, we decided to investigate the ω subunit and N terminal domain of the α subunit (α NTD) of RNAP as ADs. The ω subunit is a nonessential component of RNAP and plays a role in structurally stabilizing the holoenzyme, functionalities that can activate transcription when localized to promoter elements^35^. The ω subunit was the first AD used in a bacterial CRISPRa system in *E. coli*^18^, and has been adapted for use in other species^34^. The αNTD is responsible for initiating RNAP assembly and has recently been demonstrated as a robust AD^22^. In addition to these ADs, we also considered a transcription-factor-based AD. Specifically, we investigated the RNA polymerase binding protein A (RbpA), a transcription activator unique to Actinobacteria^36^. RbpA activates transcription through a combination of stabilizing RNAP-promoter open complex and recruiting the principal σ factor, functionalities that we hypothesized would allow RbpA to serve as an AD. Plasmids were constructed in which the αNTD, ω and RbpA ADs derived from *S. venezuelae* were translationally fused to the C terminus of dCas9. For the fusion, we employed a synthetic XTEN linker (SGSETPGTSESATPES) that has seen broad utility for creating chimeric fusions with Cas proteins^22,37,38^. To evaluate the performance of CRISPRa, we constructed a reporter cassette where an mCherry gene was placed under the control of the SP10 synthetic promoter, which we then integrated at the ΦC31 site of *S. venezuelae*. We then designed sgRNAs to direct CRISPRa to sites upstream of the SP10 promoter. Specifically, we targeted PAM sites located at 83 bp upstream of the promoter’s transcription start site (TSS) on the non-template strand and at 82 bp upstream of TSS on the template strand. As a control for non-specific activation, for each CRISPRa design, we used an off-target sgRNA binding to a non-coding region in the genome and a no-CRISPRa condition. From these measurements we observed significant activation of mCherry expression when targeting the non-template strand with the αNTD AD (Figure 2b). No activation was observed from the RbpA or the ω subunit AD, which could be due to competition with the endogenous ω subunit for the RNAP, as has been observed in other species^18,19,21,22^. Overall, these results show we can create a functional CRISPRa system for *Streptomyces* for the first time using the αNTD AD.

**Figure 2.**
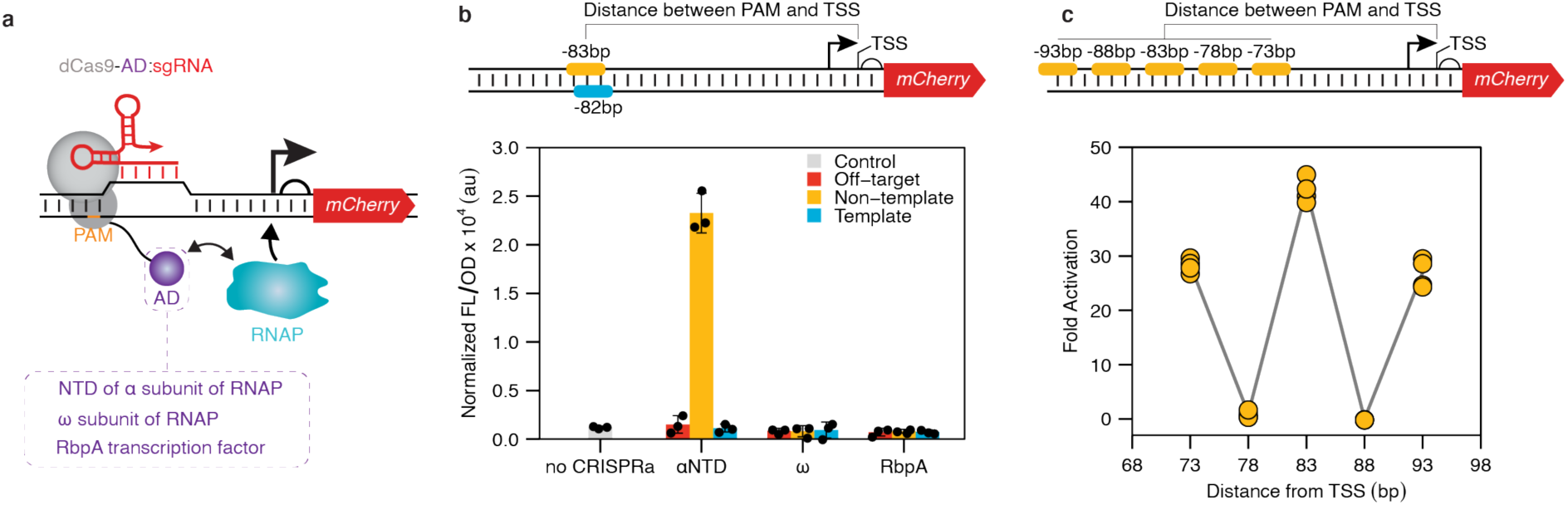
Creating a CRISPR activation (CRISPRa) for *Streptomyces venezuelae*. **a** Schematic of CRISPRa mechanism. An activator domain (AD, colored purple) is translationally fused to a dCas9 via a flexible linker. The CRISPRa complex binds upstream of a target promoter to recruit RNA Polymerase (RNAP) and activate transcription of the target gene. **b** The N-terminal domain of the α subunit of RNAP (αNTD) can serve as an AD for CRISPRa in *Streptomyces*. Schematic of sgRNA binding sites used that target the non-template (NT) and template (T) strand upstream of a promoter driving mCherry expression. The indicated distances reflect the number of nucleotides intervening between the 5’ end of the PAM (not included) and the TSS (also not included). Fluorescence characterization of *S. venezuelae* cells conjugated with CRISPRa plasmid variants using different AD. **c** CRISPRa activation shows periodical distance-dependent activation patterns. Schematic of sgRNA binding sites used that target different sites on the non-template strand upstream of a promoter driving mCherry expression. Fluorescence characterization was performed by bulk fluorescence measurements (measured in units of fluorescence [FL]/optical density [OD] at 600 nm). Fold activation was calculated by dividing the [FL]/[OD] obtained in the presence of a CRISPRa against the no-CRISPRa control within each reporter plasmid. Data are reported as individual replicates with a line connecting the mean of each condition.

### Distance dependent activation of CRISPRa in *S. venezuelae*

An intriguing design constraint of CRISPRa systems in *E. coli* are periodical activation patterns in which activation is only observed when the system is targeted to PAMs within a 2-4 bp window that repeat every 10-11 bp from the promoter’s TSS^20,22^. It has been proposed that these patterns are due to the requirement for ADs to be localized on specific faces of the DNA helix relative to the targeted promoter’s TSS^20,22^. These activation patterns have been observed for different ADs, and are independent of the dCas9-AD linker sequence and recruitment strategies being used (i.e., recruitment through binding sgRNA or dCas9)^20,22^. To determine if periodicity in activation is observed for CRISPRa in *Streptomyces*, we introduced PAM sites on the non-template strand at 73, 78, 83, 88, 93 bp upstream of the mCherry promoter’s TSS that was integrated within the *S. venezuelae* genome (Figure 2c). Corresponding sgRNAs were designed and transcription activation was measured via mCherry fluorescence. Importantly, as has been reported in *E. coli*, we observed strong activation patterns that appear to repeat with a 10 bp periodicity (Figure 2c, Supplementary Figure 4). Specifically, we see more than 20-fold activation when targeting PAMs located at 73, 83, and 93 bp from the TSS, and no activation when targeting PAMs at 78 and 88 bp. This demonstrates that CRISPRa in *Streptomyces* exhibits periodical activation patterns that need to be considered when deploying this new regulatory tool.

### Activating a silent BGC using CRISPRi and CRISPRa regulators

We next sought to demonstrate the utility of CRISPR-Cas regulatory tools for activating silent BGCs through perturbation and rewiring of endogenous regulation. For this, we decided to focus on the *S. venezuelae* jadomycin B (jdB) cluster, which encodes a type II polyketide synthase biosynthetic pathway. This cluster spans ~28 kb and includes 31 genes: 23 biosynthetic genes, 7 regulatory genes and 1 transporter gene (Figure 3a). The expression of the jdB cluster is under the control of a highly complicated gene regulatory network involving numerous regulators including JadR1, JadR2, JadR3, JadR*, and JadW1-W3. While our understanding of the regulators and network is likely incomplete, two of the most prominent regulators are JadR1 and JadR2 (Figure 3b). JadR1 is the main activator that directly turns on the expression of the *jadJ-V* operon in the presence of low levels of jdB and is essential for jdB production. However, at high levels of jdB, jadR1 represses its own promoter^39^. JadR2, a pseudo γ-butyrolactone (GBL) receptor, is the main cluster repressor as it directly shuts off jadR1 expression, thus indirectly repressing BGC activation^40^. The net result of this regulation is that under standard laboratory culturing conditions the jdB cluster is not expressed. Interestingly, in the presence of environmental stressors such as ethanol shock, heat shock, and phage infection, jadR2 repression can be relieved and jdB synthesis induced. Inspired by this, we reasoned that CRISPRi could be used to synthetically knock down the expression of JadR2, relieving JadR1 repression and thus activating jdB synthesis (Figure 3c). To test this, we created a JadR2-repressing CRISPRi plasmid and conjugated it into wild-type *S. venezuelae* cells. A no-CRISPRi condition, containing an empty plasmid, was conjugated in parallel as a negative control. After growing and fermenting the exconjugants, solvent extraction was performed on the mycelia, and liquid chromatography–mass spectrometry (LC/MS) analysis was performed on the crude extracts. From these measurements, we observed production of jadomycin B (*m/z* 550.2059, [M+H]^+^) in the presence of CRISPRi, but not in the no-CRISPRi control (Figure 3c, Supplementary Fig 5). This demonstrated that CRISPRi can be used as a tool to activate silent BGCs by relieving the regulation of endogenous repressors.

**Figure 3.**
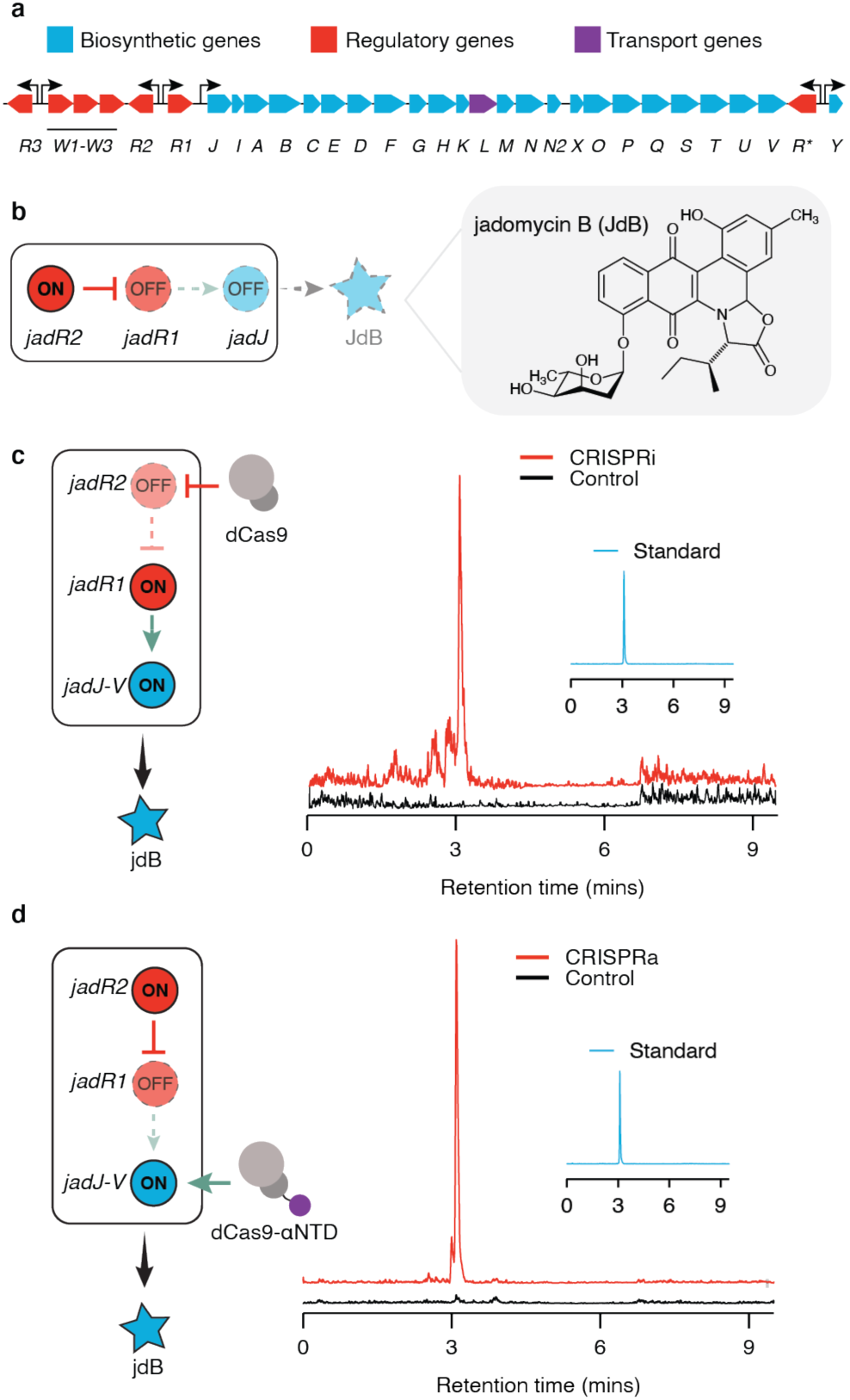
Using CRISPRi and CRISPRa to activate the silent jadomycin b (jdB) biosynthetic gene cluster (BGC). **a** Schematic of the jdB BGC of *S. venezuelae*. **b** Schematic of JadR1 and JadR2 regulation of the jdB BGC under normal laboratory conditions. Blunted arrows indicate repression and pointed arrows indicate activation. Dotted outline represents an inactive regulator and regulation step. Structure of jdB shown right of panel. **c** Schematic of CRISPRi repressing JadR1 to relieve repression on the jdB BGC and induce expression. LC-MS analysis of extracts from *S. venezuelae* conjugated with CRISPRi plasmids or no-CRISPRi control. Insert shows LC-MS analysis of a jdB standard. **d**. Schematic of CRISPRa activating the jadJ-V operon to induce expression of jdB BGC. LC-MS analysis of extracts from *S. venezuelae* conjugated with CRISPRa plasmids or no-CRISPRa control. Insert shows LC-MS analysis of a jdB standard. Data in d and e are extracted ion chromatograms at m/z 550.2059 ([M+H]+) of one representative biological replicate for each condition. Other biological replicates are shown in Supplementary figure 6.

Next, we sought to demonstrate that CRISPRa could also be used to induce activation of the jdB cluster. Specifically, our idea was to use CRISPRa to synthetically induce expression of the main biosynthetic operon (*jadJ-V*), in essence, rewiring the native regulatory network (Figure 3d). To this end, we conjugated a jadJV-activating CRISPRa plasmid into wild-type *S. venezuelae* cells. Subsequently, we tested the ability of CRISPRa to induce production of jdB by performing LC/MS analysis on the crude extracts obtained from fermentation, as described above. We detected the production of jdB in the presence of CRISPRa, but not in the control, demonstrating the ability of CRISPRa to activate silent BGCs through direct activation of key biosynthetic genes (Figure 3d, Supplementary Fig 6).

Taken together, these data show that our CRISPR regulatory tools can effectively be used to synthetically perturb and rewire the endogenous regulation of the jdB BGC and in doing so, activate the expression of this silent BGC to induce natural product synthesis.

## DISCUSSION

In our work, we establish two CRISPR-Cas systems for gene expression control in *Streptomyces*, and successfully use them to activate a silent BGC. Specifically, we optimize and resolve the design rules for our CRISPRi system in *Streptomyces* and demonstrate, for the first time, its ability to activate a silent BGC through relieving the repression of endogenous regulators. In addition, we provide the first example of a CRISPRa system for *Streptomyces*, and demonstrate the ability of this system to directly activate BGC expression through targeting ‘silent’ promoter elements. Collectively, this work provides an expanded toolbox for gene expression control in *Streptomyces* and more broadly, demonstrates a new framework for using CRISPR-Cas regulators for natural product discovery.

Our work advances the available molecular tools for engineering *Streptomyces*, which is an increasingly important genus for drug discovery and biomanufacturing. Specifically, we created an optimized CRISPRi system for *S. venezuelae* and resolved design rules. Interestingly, we observed robust repression from CRISPRi and design rules that are consistent with previous studies in bacteria^33^, adding further evidence that CRISPRi is a highly portable regulatory mechanism across the bacteria domain of life. Additionally, we created the first example of CRISPRa for *Streptomyces*. Importantly, while CRISPRi has been demonstrated across diverse bacterial species, CRISPRa systems have largely been restricted to model Gram-negative bacteria (e.g., *E. coli*), with only a single demonstration in a gram-positive bacterium, *B. subtilis*^34^. Part of the challenge of CRISPRa is the identification of functional ADs for each host. Our results, along with other recent demonstrations in *E*.*coli*^22^, suggest that host-derived αNTD AD can serve as a potentially generalizable strategy to create CRISPRa systems for diverse bacterial species. We anticipate the creation of new CRISPR-Cas regulatory systems will advance both basic science investigations of *Streptomyces* and application-specific manipulations. For example, *Streptomyces* are increasingly being utilized for commercial production of natural products^41^. However, a major challenge are low titers of natural products that require optimization to ensure viable economics^42^. Our CRISPR-Cas tools provide a new capability to program endogenous gene expression to increase flux through desired pathways, while minimizing flux through competing pathways, as has been widely demonstrated in other microbes^43–50^.

Our work introduces a novel pathway-specific approach to activate silent BGCs that complements existing strategies. Previous pathway-specific methods have largely relied on refactoring BGCs directly on the host’s genome or within heterologous hosts^11^. While successful, this approach can be arduous and challenging due to the complexity and size of BGCs. In contrast, CRISPR-Cas regulators provide a potentially more scalable framework. Libraries of sgRNAs are cheap and easy to synthesize, and regulators are easily-transferred into desired hosts through established and efficient conjugation methods. CRISPR-Cas regulators can also be used for pooled screening^51^, which we envisage could be used to multiplex different perturbations in a single experiment. More broadly, our work adds to a growing set of synthetic biology technologies for BGC activation^11,52–54^, which we anticipate can be synergistically combined in the future. For example, DNA-based transcription factor decoys have been utilized to relieve endogenous repression that could be used alongside our CRISPRi and CRISPRa technologies^53^.

While our CRISPR-Cas regulators represent a novel approach to activate BGCs, challenges remain. In particular, we observed that CRISPRa systems are only able to activate transcription when targeted to specific positions relative to the TSS. This periodical 10 bp activation pattern is proposed to be due to the rotation of the DNA double helix (~10.5 bp) and the requirement of the AD to be localized to specific faces of the DNA relative to the promoter^20,22^. While the likelihood of encountering an NGG PAM is high in *Streptomyces* spp. given their ~70% GC content^55^, not all promoters will possess optimally positioned PAMs. Importantly, several strategies have been demonstrated in *E*.*coli* to overcome this limitation, which we anticipate could be applied to *Streptomyces*. For example, engineered dCas9 proteins with relaxed PAM requirements such dxCas(3.7) have been used for CRISPRa systems to allow activation from non-canonical PAMs^20^. Additionally, circularly permuted dCas9 proteins (cpdCas9) have also been used to create CRISPRa systems with distinct, non-overlapping activation patterns^22^. Finally, alternative CRISPR-Cas systems beyond type II have also been used to create CRISPRa systems with distinct regulatory properties^56^. Beyond the limitations of CRISPRa, we anticipate more work is required to understand the generalizability of our approach. For example, it remains to be elucidated if CRISPRa is capable of activating the diversity of different endogenous promoter elements. Finally, we anticipate deeper understanding of the regulatory networks controlling natural product synthesis will be required to precisely engineer the corresponding metabolic pathways.

In summary, this work expands the current synthetic biology tools to activate silent BGCs by leveraging CRISPR-Cas regulators to perturbate and rewire the underlying regulatory gene networks. While more work is still needed to expand and generalize the approach, our system demonstrates the applicability of CRISPRi and CRISPRa systems to activate BGCs in *Streptomyces*. This method only requires the design of a specific sgRNA and the straightforward transformation of a single plasmid, thus representing a simple approach to activate silent BGCs. We anticipate that this work will pave the way for the development of high-throughput technologies for the discovery of novel secondary metabolites on a large scale.

## METHODS

### Plasmid assembly

All plasmids used in this study can be found in Supplementary Table 1 with key sequences provided in Supplementary Tables 2-5. All plasmids were constructed with either Gibson assembly^57^, Golden Gate assembly^58^, or inverse PCR (iPCR). DNA manipulations were performed in *E. coli* strain NEB Turbo. Enzymes were obtained from New England Biolabs.

### Strains and growth media

*E. coli* strains were grown in Luria Bertani (LB) broth, containing 50 μg/ml spectinomycin or 50 μg/mL apramycin as needed. 2,6-diaminopimelic acid was added to LB at a final concentration of 0.1 mM for culturing *E. coli* strain WM6029^59^. *S. venezuelae* ATCC 10712 was cultured in complete supplement medium (CSM) unless otherwise indicated. To prepare CSM, 30 g of tryptic soy broth, 1.2 g of Yeast Extract, and 1 g of MgSO_4_ were added to 1 L of water and autoclaved; filter-sterilized D-(+)-glucose and D-(+)-maltose were then added at a final concentration of 28 mM and 12 mM, respectively. Conjugation involving WM6029 and *S. venezuelae* was conducted on AS-1 medium. AS-1 was prepared by adding 5 g of soluble starch, 2.5 g of NaCl, 1 g of yeast extract and 18 g of agar to ddH_2_O to a final volume of 1 L and then autoclaved. A filter-sterilized solution of alanine, arginine and asparagine was added to a final concentration of 0.02 % w/v of each amino acid. Finally, an autoclaved Na_2_SO_4_ solution was added to a final concentration of 1 %. To prepare the MSM minimal medium for fermentation experiments, MgSO_4_ (0.04%, w/v), MOPS (0.377%, w/v), salt solution (0.9%, v/v), trace mineral solution (0.45%, v/v), and 0.2% w/v FeSO_4_·7H_2_O solution (0.45%, v/v) were added to ddH_2_O and the pH was adjusted to 7.5. The salt solution was made by addition of NaCl (1%, w/v) and CaCl2 (1%, w/v) to ddH_2_O. The trace mineral solution was made by addition of ZnSO_4_·7H_2_O (0.088%, w/v), CuSO_4_·5H_2_O (0.0039%, w/v), MnSO_4_·4H_2_O (0.00061%, w/v), H_3_BO_3_ (0.00057%, w/v), and (NH_4_)Mo_7_O_24_·4H_2_O (0.00037%, w/v) to ddH_2_O.

### Reporter strains construction

Constructs containing different promoter-mCherry combinations were assembled as described above using plasmid JBEI16292, harboring the ΦC31 integration system, as a backbone. These reporter plasmids were subsequently conjugated (see below) and integrated into *S. venezuelae* ATCC 10712, thus yielding the reporter strains.

### Interspecies conjugation

*E. coli* WM6029 cells were transformed with the plasmids to be conjugated. Colonies were picked into liquid LB media containing the appropriate antibiotic and 0.1 mM DAP, and grown overnight at 37 °C. At the same time, *S. venezuelae* mycelia were grown overnight in CSM. Liquid cultures were then pelleted and the medium was removed. Each WM6029 sample was resuspended in 500 μL of fresh LB with no antibiotics, while *S. venezuelae* pellets were resuspended in 2 mL of fresh CSM. WM6029 and *S. venezuelae* were then mixed at a 1:1 ratio, and each co-culture was spotted on AS-1 plates supplemented with 0.1 mM DAP. After incubating for 16-20 hours at 30 °C, the plates were flooded with solutions containing 500 μg of nalidixic acid and 1 mg of the appropriate antibiotic. Plates were then stored at 30 °C until the appearance of exconjugant colonies (3 to 6 days). Exconjugant colonies were streaked on fresh ISP-2 plates supplemented with either apramycin or spectinomycin at a final concentration of 50 μg/mL.

### Fluorescence measurements

*S. venezuelae* exconjugants were picked from ISP-2 plates and used to inoculate 5 mL of CSM supplemented with antibiotics as needed (apramycin or spectinomycin, final concentration 50 μg/mL). These seed cultures were grown for 2-3 days at 30 °C, then diluted to an optical density at 600 nm (OD_600_) of 0.01 in fresh CSM supplemented with antibiotics, and grown for 24 hours. After 24 hours, 25 μL of each culture were transferred to 75 μL of fresh CSM media supplemented with antibiotics as needed inside a 96 well microplate (Costar). OD_600_ (OD) and fluorescence (FL) (excitation 587 nm and emission 610 nm) were measured in a microplate reader (Tecan Infinite M1000 Pro). When performing CRISPRi and CRISPRa experiments, a *S. venezuelae* strain harboring a previously integrated mCherry reporter was used as recipient for conjugation. After conjugation, fluorescence measurement experiments were carried out as described above.

### Fluorescence data analysis

In each fluorescence measurement experiment, OD_600_ and FL values for each sample were corrected by subtracting the mean OD and FL values of a media blank. The ratio of FL to OD_600_ (FL/OD) was then calculated. Data are reported as mean FL/OD values for each condition, and error bars represent standard deviation (s.d.).

### Growth curve experiments

*S. venezuelae* colonies were grown at 30 °C with 250 rpm shaking in 5 mL CSM supplemented with apramycin or spectinomycin as needed. Upon reaching high cell density (~2 days), cells were diluted in the same medium to an OD_600_ of 0.08 directly in a Costar 96 well microplate (final volume 100 μL). Cells were then grown in a microplate reader (Tecan Spark multimode plate reader) at 30 °C with 90 rpm shaking for 24 hours. OD_600_ measurements were automatically taken every 10 minutes. For data analysis, OD_600_ values of the media controls were averaged at each time point, and then subtracted from the individual values of each condition at each time point.

### Distance-dependent CRISPRa experiments

To evaluate distance-dependent effects, sgRNAs were designed to target sequences proximal to PAMs positioned on the non-template strand at a distance of 73, 83, 93 bp upstream of the reporter promoter’s TSS. Distances do not include either the PAM or the TSS. The same sgRNAs were used to target sequences upstream of a reporter that contained 5 additional nucleotides to extend the distance between the TSS and each PAM by 5 bp (thus creating 78 and 88 bp binding sites). Fluorescence measurements were performed as described above, and FL/OD for each sample were calculated. As each reporter had slightly different mCherry expression values, data are reported as fold activation. Fold activation was calculated by normalizing FL/OD values of each sample to the mean FL/OD values of the no-CRISPRa control of each reporter. Corresponding FL/OD values used for fold activation calculations are shown in Supplementary figure 4.

### Fermentation for jadomycin B production

Fermentations were conducted under previously described conditions^60,61^. CRISPRi or CRISPRa plasmids carrying appropriate sgRNAs were conjugated into wild-type *S. venezuelae*, as described above. Exconjugants were then picked to inoculate 100 mL of CSM supplemented with apramycin, and grown to high density (usually 3-5 days) in 250 mL Erlenmeyer flasks at 30 °C. Cultures were then centrifuged, and the pellets washed twice in MSM to remove any trace of CSM. After resuspension in 6 mL of MSM, each sample was diluted in MSM supplemented with apramycin to an OD_600_ of 0.6 and a final volume of 50 mL. Fermentation cultures were incubated at 30 °C with 250 rpm shaking. After 72 hours, cultures were centrifuged and the pellets were stored at −80 °C for LC-MS analysis.

### Extraction and LC-MS analysis

Pellets obtained from fermentations were extracted with an equal volume of acetone. Mixtures were transferred to an Erlenmeyer flask and shaken at room temperature for 30 minutes at 180 rpm. Acetone was then evaporated in a rotovap at 40 °C and 200 mbar. Crude extracts were reconstituted in 5 mL of acetonitrile, and 500 μL of the reconstituted extract was combined with 500 μL of LC-MS grade water. LC-MS was carried out on an Agilent 6470B Triple Quadrupole Mass Spectrometer interface to an Agilent 1290 Infinity II HPLC system through a Jet Spray ESI source. The LC column was an Agilent 2.1 mm ID x 50 mm, 1.8 μm SB-C18 column. Mobile phase A was water with 0.1% formic acid, and mobile phase B was acetonitrile with 0.1% formic acid. LC flow rate was 0.4 ml/min. Initial conditions were 10% B up to 95% B over three minutes. The column was flushed with 95% B from 3-6 minutes and returned to initial conditions from 6-6.5 minutes. The column was equilibrated at initial conditions from 6.5-9.5 min.

### Data analysis

Experiments were performed using at least three biological replicates per condition tested unless otherwise indicated. Data for figures 1b, 1c, 1d, and 2b are reported as bars showing mean, with error bars showing standard deviation. Individual samples are also shown as black circles. Jadomycin B production experiments were performed in biological duplicates. Data in figures 3d and 3e show extracted ion chromatograms of one representative biological sample for each condition. All remaining biological replicates for LC-MS experiments are shown in Supplementary figures 5-6.

## Data availability

All source data for main figures were deposited in Rice University’s Rice Digital Scholarship Archive (DOI: https://doi.org/10.25611/5SAC-CW45)

### Acknowledgments

The authors would like to acknowledge William Metcalf (University of Illinois at Urbana-Champaign) for providing the WM6029 *E. coli* strain. The authors would like to acknowledge Jay Keasling (UC Berkeley) for providing the genomic integration plasmid JBEI16292 used to construct the reporter strains. The authors acknowledge the support of the Shared Equipment Authority at Rice University. In particular, the authors would like to acknowledge Christopher Pennington for his assistance on LC-MS experiments. This material is based upon work supported by the Welch Foundation [C-1982-20190330 to J.C] and the Alfred P. Sloan Research Fellowship [FG-2018-10500 to J.C]. J.C. is an Alfred P. Sloan Research Fellow. The authors declare no competing financial interest.

## AUTHOR CONTRIBUTIONS

A.A., M.C.V.K, K.P.C., and J.C. designed the study, A.A. performed experiments. All authors contributed to data analysis and preparation of the manuscript.

