## Supplementary material for "Activating natural product synthesis using CRISPR interference and activation systems in *Streptomyces*": Supp_Info

Andrea Ameruoso[1], Maria Claudia Villegas Kcam[1], Katherine Piper Cohen[1] and James Chappell[1,2]
1 - Department of BioSciences, Rice University, 6100 Main Street, MS 140, Houston, TX 77005, USA.
2 - Department of Bioengineering, Rice University, 6100 Main Street, MS 142, Houston, TX 77005, USA.

### TABLE OF CONTENTS

| | |
|---|---|
| **Pages 1-4** | Supplementary table 1. List of plasmids used in this study |
| **Pages 5-15** | Supplementary table 2. Example DNA plasmid sequences |
| **Page 16** | Supplementary table 3. Promoters used in this study |
| **Pages 16-17** | Supplementary table 4. sgRNA sequences used in this study |
| **Pages 18-19** | Supplementary table 5. Activator domains (ADs) used in this study |
| **Page 20** | Supplementary figure 1. Evaluating the strength of a library of Streptomyces promoters |
| **Page 21** | Supplementary figure 2. CRISPRi results in decreased fluorescence in the absence of a sgRNA or in the presence of a non-targeting sgRNA |
| **Page 22** | Supplementary figure 3. CRISPRi results in inhibition of growth in the absence of a sgRNA of in the presence of a non-targeting sgRNA |
| **Page 23** | Supplementary figure 4. Evaluating distance-dependent activation patterns of CRISPRa |
| **Page 24** | Supplementary figure 5. Activating production of jadomycin B using CRISPRi |
| **Page 25** | Supplementary figure 6. Activating production of jadomycin B using CRISPRa |
| **Page 26** | References |

**Supplementary table 1. List of all plasmids used in this study.** Abbreviations are as follows: R9 = R9 ribosome binding site, ori = origin of replication, specR = spectinomycin resistance gene, apmR = apramycin resistance gene, oriT = origin of transfer, sgRNA = single guide RNA, ds origin = double-stranded origin, bp = base pairs, NT = non-template strand, T = template strand, αNTD = N-terminal domain of the α subunit of RNAP. Promoters: KasO*p, ermE*p, gapdh(EL), rpsL(XC), 57, SP43, SP30, SP20, SP10, SP1. Terminators: T7, λt0, Fd. Origins of replication: pMB1, pUC.

| Plasmid ID | Plasmid features | Name | Figure(s) |
|---|---|---|---|
| pJEC532 | KasO*p - mCherry - T7 - pMB1 ori - SpecR - RP4 oriT - ΦC31 attP site - ΦC31 integrase | mCherry reporter | 1b, 1c, 1d, S1 |
| pJEC533 | gapdh(EL) - mCherry - T7 terminator - pMB1 ori - SpecR - RP4 oriT - ΦC31 attP site - ΦC31 integrase | gapdh(EL)-mCherry | S1 |
| pJEC710 | Fd terminator - gapdh(EL) - lacZ - sgRNA scaffold - λt0 terminator - pUC ori - ApmR - pSG5 replicase - pSG5 ds origin - RP4 oriT | no CRISPR | 1b, 1c, 1d, 2b, 2c, 3d, 3e, S1, S2, S3 |
| pJEC711 | rpsL(XC) - dCas9 - Fd terminator - gapdh(EL) - mCherry sgRNA (+11bp) - sgRNA scaffold - λt0 terminator - pUC ori - ApmR - pSG5 replicase - pSG5 ds origin - RP4 oriT | rpsL(XC)-dCas9/gapdh(EL)-sgRNA | 1b |
| pJEC712 | rpsL(XC) - dCas9 - Fd terminator - SP43 - mCherry sgRNA (+11bp) - sgRNA scaffold - λt0 terminator - pUC ori - ApmR - pSG5 replicase - pSG5 ds origin - RP4 oriT | rpsL(XC)-dCas9/SP43-sgRNA | 1b, 1c |
| pJEC713 | SP1 - RiboJ - R9 - dCas9 - Fd terminator - SP43 - mCherry sgRNA (+11bp) - sgRNA scaffold - λt0 terminator - pUC ori - ApmR - pSG5 replicase - pSG5 ds origin - RP4 oriT | SP1-dCas9/SP43-sgRNA | 1c |
| pJEC714 | SP30 - RiboJ -R9 - dCas9 - Fd terminator - SP43 - mCherry sgRNA (+11bp NT) - sgRNA scaffold - λt0 terminator - pUC ori - ApmR - pSG5 replicase - pSG5 ds origin - RP4 oriT | SP30-dCas9/SP43-sgRNA (+11 NT) | 1c, 1d, S2 |
| pJEC715 | SP43 - RiboJ -R9 - mCherry - T7 terminator - pMB1 ori - SpecR - RP4 oriT - ΦC31 attP site - ΦC31 integrase | SP43-mCherry | S1 |
| pJEC716 | SP30 - RiboJ -R9 - mCherry - T7 terminator - pMB1 ori - SpecR - RP4 oriT - ΦC31 attP site - ΦC31 integrase | SP30-mCherry | S1 |
| pJEC717 | ermE*p - mCherry - T7 terminator - pMB1 ori - SpecR - RP4 oriT - ΦC31 attP site - ΦC31 integrase | ermE*p-mCherry | S1 |
| pJEC718 | SP20 - RiboJ -R9 - mCherry - T7 terminator - pMB1 ori - SpecR - RP4 oriT - ΦC31 attP site - ΦC31 integrase | SP20-mCherry | S1 |
| pJEC719 | SP10 - RiboJ -R9 - mCherry - T7 terminator - pMB1 ori - SpecR - RP4 oriT - ΦC31 attP site - ΦC31 integrase | SP10-mCherry | 2b, 2c, S1 |

| | | | |
|---|---|---|---|
| pJEC720 | rpsL(XC) - mCherry - T7 terminator - pMB1 ori - SpecR - RP4 oriT - ΦC31 attP site - ΦC31 integrase | rpsL(XC)-mCherry | S1 |
| pJEC721 | 57 - mCherry - T7 terminator - pMB1 ori - SpecR - RP4 oriT - ΦC31 attP site - ΦC31 integrase | 57-mCherry | S1 |
| pJEC722 | SP1 - RiboJ -R9 - mCherry - T7 terminator - pMB1 ori - SpecR - RP4 oriT - ΦC31 attP site - ΦC31 integrase | SP1-mCherry | S1 |
| pJEC723 | SP30 - RiboJ -R9 - dCas9 - Fd terminator - SP43 - non-coding genomic region sgRNA #1 - sgRNA scaffold - λt0 terminator - pUC ori - ApmR - pSG5 replicase - pSG5 ds origin - RP4 oriT | CRISPRi with sgRNA binding to a genomic region | 1d,S2 |
| pJEC724 | SP30 - RiboJ -R9 - dCas9 - Fd terminator - SP43 - non-coding genomic region sgRNA #2 - sgRNA scaffold - λt0 terminator - pUC ori - ApmR - pSG5 replicase - pSG5 ds origin - RP4 oriT | CRISPRi with sgRNA binding to a genomic region | S2 |
| pJEC725 | SP30 - RiboJ -R9 - dCas9 - Fd terminator - SP43 - no-match sgRNA #2 - sgRNA scaffold - λt0 terminator - pUC ori - ApmR - pSG5 replicase - pSG5 ds origin - RP4 oriT | SP30-dCas9/SP43-sgRNA off-target w/o binding site | S2 |
| pJEC726 | SP30 - RiboJ -R9 - dCas9 - Fd terminator - SP43 - lacZ - sgRNA scaffold - λt0 terminator - pUC ori - ApmR - pSG5 replicase - pSG5 ds origin - RP4 oriT | SP30-dCas9/SP43-sgRNA scaffold (i.e. no sgRNA control) | S2 |
| pJEC727 | SP30 - RiboJ -R9 - dCas9 - Fd terminator - SP43 - mCherry sgRNA (+123bp NT) - sgRNA scaffold - λt0 terminator - pUC ori - ApmR - pSG5 replicase - pSG5 ds origin - RP4 oriT | SP30-dCas9/SP43-sgRNA (+123 NT) | 1d |
| pJEC728 | SP30 - RiboJ -R9 - dCas9 - Fd terminator - SP43 - mCherry sgRNA (+230bp NT) - sgRNA scaffold - λt0 terminator - pUC ori - ApmR - pSG5 replicase - pSG5 ds origin - RP4 oriT | SP30-dCas9/SP43-sgRNA (+230 NT) | 1d |
| pJEC729 | SP30 - RiboJ -R9 - dCas9 - Fd terminator - SP43 - mCherry sgRNA (+531bp NT) - sgRNA scaffold - λt0 terminator - pUC ori - ApmR - pSG5 replicase - pSG5 ds origin - RP4 oriT | SP30-dCas9/SP43-sgRNA (+531 NT) | 1d |
| pJEC730 | SP30 - RiboJ -R9 - dCas9 - Fd terminator - SP43 - mCherry sgRNA (+623bp NT) - sgRNA scaffold - λt0 terminator - pUC ori - ApmR - pSG5 replicase - pSG5 ds origin - RP4 oriT | SP30-dCas9/SP43-sgRNA (+623 NT) | 1d |
| pJEC731 | SP30 - RiboJ -R9 - dCas9 - Fd terminator - SP43 - mCherry sgRNA (+9bp T) - sgRNA scaffold - λt0 terminator - pUC ori - ApmR - pSG5 replicase - pSG5 ds origin - RP4 oriT | SP30-dCas9/SP43-sgRNA (+9 T) | 1d |

| | | | |
|---|---|---|---|
| pJEC732 | SP30 - RiboJ -R9 - dCas9 - Fd terminator - SP43 - mCherry sgRNA (+112bp T) - sgRNA scaffold - λt0 terminator - pUC ori - ApmR - pSG5 replicase - pSG5 ds origin - RP4 oriT | SP30-dCas9/SP43-sgRNA (+112 T) | 1d |
| pJEC733 | SP30 - RiboJ -R9 - dCas9 - Fd terminator - SP43 - mCherry sgRNA (+223bp T) - sgRNA scaffold - λt0 terminator - pUC ori - ApmR - pSG5 replicase - pSG5 ds origin - RP4 oriT | SP30-dCas9/SP43-sgRNA (+223 T) | 1d |
| pJEC734 | SP30 - RiboJ -R9 - dCas9 - Fd terminator - SP43 - mCherry sgRNA (+540bp T) - sgRNA scaffold - λt0 terminator - pUC ori - ApmR - pSG5 replicase - pSG5 ds origin - RP4 oriT | SP30-dCas9/SP43-sgRNA (+540 T) | 1d |
| pJEC735 | SP30 - RiboJ -R9 - dCas9 - Fd terminator - SP43 - mCherry sgRNA (+627bp T) - sgRNA scaffold - λt0 terminator - pUC ori - ApmR - pSG5 replicase - pSG5 ds origin - RP4 oriT | SP30-dCas9/SP43-sgRNA (+627 T) | 1d |
| pJEC736 | PAM region - SP10 - RiboJ -R9 - mCherry - T7 terminator - pMB1 ori - SpecR - RP4 oriT - ΦC31 attP site - ΦC31 integrase | CRISPRa reporter #1 | 2b, 2c, S3 |
| pJEC737 | SP30 - RiboJ -R9 - dCas9 - XTEN - αNTD - Fd terminator - SP43 - non-coding genomic region sgRNA #1 - sgRNA scaffold - λt0 terminator - pUC ori - ApmR - pSG5 replicase - pSG5 ds origin - RP4 oriT | SP30-dCas9-αNTD/off-target | 2b |
| pJEC738 | SP30 - RiboJ -R9 - dCas9 - XTEN - αNTD - Fd terminator - SP43 - CRISPRa sgRNA (82bp T) - sgRNA scaffold - λt0 terminator - pUC ori - ApmR - pSG5 replicase - pSG5 ds origin - RP4 oriT | SP30-dCas9-αNTD/sgRNA - 82 T | 2b |
| pJEC739 | SP30 - RiboJ -R9 - dCas9 - XTEN - αNTD - Fd terminator - SP43 - CRISPRa sgRNA (83bp NT) - sgRNA scaffold - λt0 terminator - pUC ori - ApmR - pSG5 replicase - pSG5 ds origin - RP4 oriT | SP30-dCas9-αNTD/sgRNA - 83 NT | 2b, 2c, S3 |
| pJEC740 | SP30 - RiboJ -R9 - dCas9 - XTEN - ω - Fd terminator - SP43 - non-coding genomic region sgRNA #1 - sgRNA scaffold - λt0 terminator - pUC ori - ApmR - pSG5 replicase - pSG5 ds origin - RP4 oriT | SP30-dCas9-ω/off-target | 2b |
| pJEC741 | SP30 - RiboJ -R9 - dCas9 - XTEN - ω - Fd terminator - SP43 - CRISPRa sgRNA (83bp NT) - sgRNA scaffold - λt0 terminator - pUC ori - ApmR - pSG5 replicase - pSG5 ds origin - RP4 oriT | SP30-dCas9-ω/sgRNA -83 NT | 2b |
| pJEC742 | SP30 - RiboJ -R9 - dCas9 - XTEN - ω - Fd terminator - SP43 - CRISPRa sgRNA (82bp T) - sgRNA scaffold - λt0 terminator - pUC ori - ApmR - pSG5 replicase - pSG5 ds origin - RP4 oriT | SP30-dCas9-ω/sgRNA -82 T | 2b |

| | | | |
|---|---|---|---|
| pJEC743 | SP30 - RiboJ -R9 - dCas9 - XTEN - RbpA - Fd terminator - SP43 - non-coding genomic region sgRNA #1 - sgRNA scaffold - λt0 terminator - pUC ori - ApmR - pSG5 replicase - pSG5 ds origin - RP4 oriT | SP30-dCas9-RbpA/off-target | 2b |
| pJEC744 | SP30 - RiboJ -R9 - dCas9 - XTEN - RbpA - Fd terminator - SP43 - CRISPRa sgRNA (83bp NT) - sgRNA scaffold - λt0 terminator - pUC ori - ApmR - pSG5 replicase - pSG5 ds origin - RP4 oriT | SP30-dCas9-RbpA/sgRNA - 83 NT | 2b |
| pJEC745 | SP30 - RiboJ -R9 - dCas9 - XTEN - RbpA - Fd terminator - SP43 - CRISPRa sgRNA (82bp T) - sgRNA scaffold - λt0 terminator - pUC ori - ApmR - pSG5 replicase - pSG5 ds origin - RP4 oriT | SP30-dCas9-RbpA/sgRNA - 82 T | 2b |
| pJEC746 | PAM region + 5bp - SP10 - RiboJ -R9 - mCherry - T7 terminator - pMB1 ori - SpecR - RP4 oriT - ΦC31 attP site - ΦC31 integrase | CRISPRa reporter #2 (PAMs shifted by 5bp) | 2c, S3 |
| pJEC747 | SP30 - RiboJ -R9 - dCas9 - XTEN - αNTD - Fd terminator - SP43 - CRISPRa sgRNA (73bp NT) - sgRNA scaffold - λt0 terminator - pUC ori - ApmR - pSG5 replicase - pSG5 ds origin - RP4 oriT | SP30-dCas9-αNTD/sgRNA - 73 NT | 2c, S3 |
| pJEC748 | SP30 - RiboJ -R9 - dCas9 - XTEN - αNTD - Fd terminator - SP43 - CRISPRa sgRNA (93bp NT) - sgRNA scaffold - λt0 terminator - pUC ori - ApmR - pSG5 replicase - pSG5 ds origin - RP4 oriT | SP30-dCas9-αNTD/sgRNA - 93 NT | 2c, S3 |
| pJEC749 | SP30 - RiboJ -R9 - dCas9 - Fd terminator - SP43 - CRISPRi jadR2 sgRNA (+44bp NT) - sgRNA scaffold - λt0 terminator - pUC ori - ApmR - pSG5 replicase - pSG5 ds origin - RP4 oriT | jadR2 CRISPRi | 3d |
| pJEC750 | SP30 - RiboJ -R9 - dCas9 - XTEN - αNTD - Fd terminator - SP43 - CRISPRa jadJ sgRNA (-73bp NT) - sgRNA scaffold - λt0 terminator - pUC ori - ApmR - pSG5 replicase - pSG5 ds origin - RP4 oriT | jadJ CRISPRa | 3e |

**Supplementary table 2. Example DNA plasmid sequences.**

| Name and features | DNA sequence |
|---|---|
| CRISPRi reporter<br><br>(Promoter-RBS-mCherry-Terminator-E.coli ori-SpecR-oriT-attP site-phiC31 integrase) | CGAGACACCCGGGAAGCCTGATCTACGTCTGTCGAGAAGTTTCTGATCG<br>ATGACACTCGTTCGTTCACACGTTGCAGCAGAGTACTTGTTCACATTCGA<br>ACGGTCTCTGCTTTGACAACATGCTGTGCGGTGTTGTAAAGTCGTGGCC<br>AGGAGAATACGACAGCGTGCAGGACTGGGGGAGTGCGCATATGGTCTC<br>CAAGGGCGAGGAGGACAACATGGCCATCATCAAGGAGTTCATGCGCTTC<br>AAGGTCCACATGGAGGGCTCCGTCAACGGGCACGAGTTCGAGATCGAG<br>GGCGAGGGGGAGGGCCGGCCGTACGAGGGCACCCAGACCGCCAAGCT<br>GAAGGTGACCAAGGGCGGCCCCCTCCCGTTCGCCTGGGACATCCTCTC<br>CCCCCAGTTCATGTACGGCTCGAAGGCCTACGTCAAGCACCCGGCCGA<br>CATCCCGGACTACCTGAAGCTCTCGTTCCCGGAGGGGTTCAAGTGGGA<br>GCGGGTCATGAACTTCGAGGACGGCGGCGTCGTCACCGTCACCCAGGA<br>CAGCTCCCTGCAGGACGGCGAGTTCATCTACAAGGTCAAGCTGCGGGG<br>CACGAACTTCCCGAGCGACGGCCCCGTGATGCAGAAGAAGACGATGGG<br>CTGGGAAGCGTCCTCGGAGCGCATGTACCCGGAGGACGGCGCCCTCAA<br>GGGCGAGATCAAGCAGCGCCTGAAGCTGAAGGACGGCGGCCACTACGA<br>CGCCGAAGTCAAGACGACGTACAAGGCCAAGAAGCCGGTGCAGCTCCC<br>GGACGCCTACAACGTGAACATCAAGCTCGACATCACCTCGCACAACGAG<br>GACTACACGATCGTGGAGCAGTACGAGCGCGCCGAGGGCCGGCACTCG<br>ACCGGCGGCATGGACGAGCTGTACAAGTGAGCCGTTCGCGCCGCCCCG<br>GCTCGCATCGTCCCCGACCCGTCAACCTCATCCGCAAGGAGTCTCTAGA<br>GGATCCGCGGCCGCGCGATATCGAATTCCTCGAGTAACTAGCATAAC<br>CCCTTGGGGCCTCTAAACGGGTCTTGAGGGGTTTTTTGTTAAGCCGAAC<br>AGGAAGCACAGCTCCTACTGAACATGTGAGCAAAAGGCCAGCAAAAGGC<br>CAGGAACCGTAAAAAGGCCGCGTTGCTGGCGTTTTTCCATAGGCTCCGC<br>CCCCCTGACGAGCATCACAAAAATCGACGCTCAAGTCAGAGGTGGCGAA<br>ACCCGACAGGACTATAAAGATACCAGGCGTTTCCCCCTGGAAGCTCCCT<br>CGTGCGCTCTCCTGTTCCGACCCTGCCGCTTACCGGATACCTGTCCGCC<br>TTTCTCCCTTCGGGAAGCGTGGCGCTTTCTCATAGCTCACGCTGTAGGT<br>ATCTCAGTTCGGTGTAGGTCGTTCGCTCCAAGCTGGGCTGTGTGCACGA<br>ACCCCCCGTTCAGCCCGACCGCTGCGCCTTATCCGGTAACTATCGTCTT<br>GAGTCCAACCCGGTAAGACACGACTTATCGCCACTGGCAGCAGCCACTG<br>GTAACAGGATTAGCAGAGCGAGGTATGTAGGCGGTGCTACAGAGTTCTT<br>GAAGTGGTGGCCTAACTACGGCTACACTAGAAGAACAGTATTTGGTATCT<br>GCGCTCTGCTGAAGCCAGTTACCTTCGGAAAAAGAGTTGGTAGCTCTTG<br>ATCCGGCAAACAAACCACCGCTGGTAGCGGTGGTTTTTTTGTTTGCAAG<br>CAGCAGATTACGCGCAGAAAAAAAGGATCTCAAGAAGATCCTTTGATCTT<br>TTCTACGGGGTCTGACGCTCAGTGGAACGAAAACTCACGTTAAGGGATT<br>TTGGTCATGAGATTATCAAAAAGGATCTTCACCTAGATCCTTTTGGTTCAT<br>GTGCAGCTCCATCAGCAAAAGGGGATGATAAGTTTATCACCACCGACTAT<br>TTGCAACAGTGCCGTTGATCGTGCTATGATCGACTGATGTCATCAGCGG<br>TGGAGTGCAATGTCATGCGCTCACGCAACTGGTCCAGAACCTTGACCGA<br>ACGCAGCGGTGGTAACGGCGCAGTGGCGGTTTTCATGGCTTGTTATGAC<br>TGTTTTTTTGGGGTACAGTCTATGCCTCGGGCATCCAAGCAGCAAGCGC<br>GTTACGCCGTGGGTCGATGTTTGATGTTATGGAGCAGCAACGATGTTAC<br>GCAGCAGGGCAGTCGCCCTAAAACAAAGTTAAACATC<br>ATGAGGGAAGCGGTGATCGCCGAAGTATCGACTCAACTATCAGAGGTAG<br>TTGGCGTCATCGAGCGCCATCTCGAACCGACGTTGCTGGCCGTACATTT |

GTACGGCTCCGCAGTGGATGGCGGCCTGAAGCCACACAGTGATATTGAT
TTGCTGGTTACGGTGACCGTAAGGCTTGATGAAACAACGCGGCGAGCTT
TGATCAACGACCTTTTGGAAACTTCGGCTTCCCCTGGAGAGAGCGAGAT
TCTCCGCGCTGTAGAAGTCACCATTGTTGTGCACGACGACATCATTCCGT
GGCGTTATCCAGCTAAGCGCGAACTGCAATTTGGAGAATGGCAGCGCAA
TGACATTCTTGCAGGTATCTTCGAGCCAGCCACGATCGACATTGATCTGG
CTATCTTGCTGACAAAAGCAAGAGAACATAGCGTTGCCTTGGTAGGTCCA
GCGGCGGAGGAACTCTTTGATCCGGTTCCTGAACAGGATCTATTTGAGG
CGCTAAATGAAACCTTAACGCTATGGAACTCGCCGCCCGACTGGGCTGG
CGATGAGCGAAATGTAGTGCTTACGTTGTCCCGCATTTGGTACAGCGCA
GTAACCGGCAAAATCGCGCCGAAGGATGTCGCTGCCGACTGGGCAATG
GAGCGCCTGCCGGCCCAGTATCAGCCCGTCATACTTGAAGCTAGACAG
GCTTATCTTGGACAAGAAGAAGATCGCTTGGCCTCGCGCGCAGATCAGT
TGGAAGAATTTGTCCACTACGTGAAAGGCGAGATCACCAAGGTAGTCGG
CAAATAATTGTCTTTCTTCAGCTCGCTGATGATATGCCTTCCTGGTTGGC
TTGGTTTCATCAGCCATCCGCTTGCCCTCATCTGTTACGCCGGCGGTAG
CCGGCCAGCCTCGCAGAGCAGGATTCCCGTTGAGCACCGCCAGGTGCG
AATAAGGGACAGTGAAGAAGGAACACCCGCTCGCGGGTGGCCTACTT
CACCTATCCTGCCCGGCTGACGCCGTTGGATACACCAAGGAAAGTCTAC
ACGAACCCTTTGGCAAAATCCTGTATATCGTGCGAAAAAGGATGGATATA
CCGAAAAAATCGCTATAATGACCCCGAAGCAGGGTTATGCAGCGGAAAA
GATCCGTCGACCTGCAGGCATGCAAGCTCTAGCGATTCCAGACGTCCCG
AAGGCGTGGCGCGGCTTCCCCGTGCCGGAGCAATCGCCCTGGGTGGGT
TACACGACGCCCCTCTATGGCCCGTACTGACGGACACACCGAAGCCCC
GGCGGCAACCCTCAGCGGATGCCCCGGGGCTTCACGTTTTCCCAGGTC
AGAAGCGGTTTTCGGGAGTAGTGCCCCAACTGGGGTAACCTTTGAGTTC
TCTCAGTTGGGGGCGTAGGGTCGCCGACATGACACAAGGGGTTGTGAC
CGGGGTGGACACGTACGCGGGTGCTTACGACCGTCAGTCGCGCGAGCG
CGAGAATTCGAGCGCAGCAAGCCCAGCGACACAGCGTAGCGCCAACGA
AGACAAGGCGGCCGACCTTCAGCGCGAAGTCGAGCGCGACGGGGGCC
GGTTCAGGTTCGTCGGGCATTTCAGCGAAGCGCCGGGCACGTCGGCGT
TCGGGACGGCGGAGCGCCCGGAGTTCGAACGCATCCTGAACGAATGCC
GCGCCGGGCGGCTCAACATGATCATTGTCTATGACGTGTCGCGCTTCTC
GCGCCTGAAGGTCATGGACGCGATTCCGATTGTCTCGGAATTGCTCGCC
CTGGGCGTGACGATTGTTTCCACTCAGGAAGGCGTCTTCCGGCAGGGAA
ACGTCATGGACCTGATTCACCTGATTATGCGGCTCGACGCGTCGCACAA
AGAATCTTCGCTGAAGTCGGCGAAGATTCTCGACACGAAGAACCTTCAG
CGCGAATTGGGCGGGTACGTCGGCGGGAAGGCGCCTTACGGCTTCGAG
CTTGTTTCGGAGACGAAGGAGATCACGCGCAACGGCCGAATGGTCAATG
TCGTCATCAACAAGCTTGCGCACTCGACCACTCCCCTTACCGGACCCTT
CGAGTTCGAGCCCGACGTAATCCGGTGGTGGTGGCGTGAGATCAAGAC
GCACAAACACCTTCCCTTCAAGCCGGGCAGTCAAGCCGCCATTCACCCG
GGCAGCATCACGGGGCTTTGTAAGCGCATGGACGCTGACGCCGTGCCG
ACCCGGGGCGAGACGATTGGGAAGAAGACCGCTTCAAGCGCCTGGGAC
CCGGCAACCGTTATGCGAATCCTTCGGGACCCGCGTATTGCGGGCTTCG
CCGCTGAGGTGATCTACAAGAAGAAGCCGGACGGCACGCCGACCACGA
AGATTGAGGGTTACCGCATTCAGCGCGACCCGATCACGCTCCGGCCGG
TCGAGCTTGATTGCGGACCGATCATCGAGCCCGCTGAGTGGTATGAGCT
TCAGGCGTGGTTGGACGGCAGGGGGCGCGGCAAGGGGCTTTCCCGGG
GGCAAGCCATTCTGTCCGCCATGGACAAGCTGTACTGCGAGTGTGGCG
CCGTCATGACTTCGAAGCGCGGGGAAGAATCGATCAAGGACTCTTACCG

| | |
|---|---|
| | CTGCCGTCGCCGGAAGGTGGTCGACCCGTCCGCACCTGGGCAGCACGA<br>AGGCACGTGCAACGTCAGCATGGCGGCACTCGACAAGTTCGTTGCGGA<br>ACGCATCTTCAACAAGATCAGGCACGCCGAAGGCGACGAAGAGACGTTG<br>GCGCTTCTGTGGGAAGCCGCCCGACGCTTCGGCAAGCTCACTGAGGCG<br>CCTGAGAAGAGCGGCGAACGGGCGAACCTTGTTGCGGAGCGCGCCGAC<br>GCCCTGAACGCCCTTGAAGAGCTGTACGAAGACCGCGCGGCAGGCGCG<br>TACGACGGACCCGTTGGCAGGAAGCACTTCCGGAAGCAACAGGCAGCG<br>CTGACGCTCCGGCAGCAAGGGGCGGAAGAGCGGCTTGCCGAACTTGAA<br>GCCGCCGAAGCCCCGAAGCTTCCCCTTGACCAATGGTTCCCCGAAGAC<br>GCCGACGCTGACCCGACCGGCCCTAAGTCGTGGTGGGGGCGCGCGTC<br>AGTAGACGACAAGCGCGTGTTCGTCGGGCTCTTCGTAGACAAGATCGTT<br>GTCACGAAGTCGACTACGGGCAGGGGGCAGGGAACGCCCATCGAGAAG<br>CGCGCTTCGATCACGTGGGCGAAGCCGCCGACCGACGACGACGAAGAC<br>GACGCCCAGGACGGCACGGAAGACGTAGCGGCGTAG |
| CRISPRi<br><br>(Promoter-RiboJ-RBS-dCas9-terminator-promoter-sgRNA targeting sequence-sgRNA scaffold-terminator-ori-AprR-pSG5 rep-pSG5 ds-oriT) | TGTTCACATTCGAACCGTCTCTGCTTTGACATCGTGTGGCGCTTGGGTGT<br>AAAGTCGTGGCCATTAAACAAAATTATTTGTAGAGGCTGTTTCGTCCTCA<br>CGGACTCATCAGACCGGAAAGCACATCCGGTGACAGCTAACTACGAAGG<br>GGAGTCAGTATGGACAAGAAGTACAGCATCGGCCTGGCCATCGGCACC<br>AACAGCGTGGGCTGGGCGGTCATCACCGACGAGTACAAGGTCCCCTCC<br>AAGAAGTTCAAGGTCCTGGGCAACACCGACCGGCACTCGATCAAGAAGA<br>ACCTGATCGGCGCCCTGCTCTTCGACAGCGGCGAAACCGCCGAGGCGA<br>CCCGCCTGAAGCGGACCGCCCGTCGCCGCTACACCCGGCGCAAGAACC<br>GCATCTGCTACCTGCAGGAGATCTTCTCCAACGAGATGGCCAAGGTCGA<br>CGACTCGTTCTTCCACCGGCTCGAGGAGAGCTTCCTGGTGGAGGAGGA<br>CAAGAAGCACGAGCGCCACCCGATCTTCGGCAACATCGTCGACGAGGT<br>GGCCTACCACGAGAAGTACCCCACCATCTACCACCTCCGCAAGAAGCTG<br>GTCGACTCGACCGACAAGGCGGACCTGCGGCTCATCTACCTGGCCCTC<br>GCGCACATGATCAAGTTCCGCGGCCACTTCCTCATCGAGGGCGACCTGA<br>ACCCGGACAACTCCGACGTCGACAAGCTCTTCATCCAGCTGGTGCAGAC<br>CTACAACCAGCTGTTCGAGGAGAACCCCATCAACGCCAGCGGCGTCGA<br>CGCCAAGGCGATCCTCTCCGCGCGCCTGAGCAAGTCCCGGCGCCTGGA<br>GAACCTCATCGCCCAGCTGCCGGGCGAGAAGAAGAACGGCCTCTTCGG<br>CAACCTGATCGCGCTGTCGCTCGGCCTGACCCCCAACTTCAAGAGCAAC<br>TTCGACCTGGCCGAGGACGCGAAGCTCCAGCTGTCCAAGGACACCTAC<br>GACGACGACCTGGACAACCTGCTCGCCCAGATCGGCGACCAGTACGCG<br>GACCTCTTCCTGGCCGCGAAGAACCTCTCGGACGCCATCCTGCTCAGCG<br>ACATCCTGCGGGTCAACACCGAGATCACCAAGGCCCCGCTGTCGGCGA<br>GCATGATCAAGCGGTACGACGAGCACCACCAGGACCTGACCCTGCTCAA<br>GGCCCTCGTGCGCCAGCAGCTGCCCGAGAAGTACAAGGAGATCTTCTTC<br>GACCAGTCCAAGAACGGCTACGCCGGCTACATCGACGGCGGCGCGTCG<br>CAGGAGGAGTTCTACAAGTTCATCAAGCCGATCCTGGAGAAGATGGACG<br>GCACCGAGGAGCTGCTCGTCAAGCTGAACCGCGAGGACCTGCTCCGCA<br>AGCAGCGGACCTTCGACAACGGCTCCATCCCGCACCAGATCCACCTGG<br>GCGAGCTCCACGCCATCCTCCGGCGCCAGGAGGACTTCTACCCCTTCCT<br>GAAGGACAACCGCGAGAAGATCGAGAAGATCCTGACCTTCCGGATCCC<br>GTACTACGTCGGCCCCCTGGCCCGCGGCAACTCCCGGTTCGCGTGGAT<br>GACCCGGAAGTCGGAGGAAACCATCACCCCGTGGAACTTCGAGGAGGT<br>CGTGGACAAGGGCGCCTCCGCGCAGTCGTTCATCGAGCGCATGACCAA<br>CTTCGACAAGAACCTCCCGAACGAGAAGGTCCTGCCCAAGCACAGCCTG<br>CTCTACGAGTACTTCACCGTGTACAACGAGCTGACCAAGGTCAAGTACG<br>TGACCGAGGGCATGCGGAAGCCGGCCTTCCTGTCCGGCGAGCAGAAGA |

| | |
|---|---|
| | AGGCGATCGTCGACCTGCTCTTCAAGACCAACCGCAAGGTCACCGTGAA<br>GCAGCTGAAGGAGGACTACTTCAAGAAGATCGAGTGCTTCGACTCCGTC<br>GAGATCTCGGGCGTGGAGGACCGCTTCAACGCCTCCCTGGGCACCTAC<br>CACGACCTGCTCAAGATCATCAAGGACAAGGACTTCCTCGACAACGAGG<br>AGAACGAGGACATCCTGGAGGACATCGTCCTCACCCTGACCCTCTTCGA<br>GGACCGCGAGATGATCGAGGAGCGGCTCAAGACCTACGCCCACCTGTT<br>CGACGACAAGGTGATGAAGCAGCTGAAGCGGCGCCGGTACACCGGCTG<br>GGGCCGCCTCTCCCGGAAGCTGATCAACGGCATCCGGGACAAGCAGAG<br>CGGCAAGACCATCCTGGACTTCCTCAAGTCCGACGGCTTCGCCAACCGC<br>AACTTCATGCAGCTCATCCACGACGACTCGCTGACCTTCAAGGAGGACA<br>TCCAGAAGGCCCAGGTGTCCGGCCAGGGCGACAGCCTCCACGAGCACA<br>TCGCCAACCTGGCGGGCTCCCCGGCGATCAAGAAGGGCATCCTCCAGA<br>CCGTCAAGGTCGTGGACGAGCTGGTCAAGGTGATGGGCCGCCACAAGC<br>CCGAGAACATCGTGATCGAGATGGCCCGGGAGAACCAGACCACCCAGA<br>AGGGCCAGAAGAACTCCCGCGAGCGGATGAAGCGCATCGAGGAGGGCA<br>TCAAGGAGCTCGGCTCGCAGATCCTGAAGGAGCACCCGGTCGAGAACA<br>CCCAGCTCCAGAACGAGAAGCTGTACCTCTACTACCTGCAGAACGGCCG<br>CGACATGTACGTGGACCAGGAGCTCGACATCAACCGGCTGAGCGACTA<br>CGACGTCGACGCCATCGTGCCGCAGTCCTTCCTGAAGGACGACTCGATC<br>GACAACAAGGTCCTGACCCGCTCCGACAAGAACCGGGGCAAGTCCGAC<br>AACGTGCCCTCGGAGGAGGTCGTGAAGAAGATGAAGAACTACTGGCGC<br>CAGCTGCTCAACGCCAAGCTCATCACCCAGCGCAAGTTCGACAACCTGA<br>CCAAGGCCGAGCGGGGCGGCCTGTCGGAGCTCGACAAGGCGGGCTTC<br>ATCAAGCGCCAGCTCGTCGAAACCCGGCAGATCACCAAGCACGTGGCC<br>CAGATCCTGGACAGCCGGATGAACACCAAGTACGACGAGAACGACAAG<br>CTGATCCGCGAGGTCAAGGTGATCACCCTCAAGAGCAAGCTGGTGTCCG<br>ACTTCCGCAAGGACTTCCAGTTCTACAAGGTCCGGGAGATCAACAACTA<br>CCACCACGCCCACGACGCGTACCTGAACGCCGTCGTGGGCACCGCGCT<br>GATCAAGAAGTACCCGAAGCTGGAGTCCGAGTTCGTCTACGGCGACTAC<br>AAGGTCTACGACGTGCGCAAGATGATCGCCAAGTCGGAGCAGGAGATC<br>GGCAAGGCCACCGCGAAGTACTTCTTCTACAGCAACATCATGAACTTCTT<br>CAAGACCGAGATCACCCTGGCCAACGGCGAGATCCGCAAGCGGCCCCT<br>GATCGAAACCAACGGCGAAACCGGCGAGATCGTCTGGGACAAGGGCCG<br>CGACTTCGCCACCGTCCGGAAGGTGCTGTCCATGCCGCAGGTCAACATC<br>GTCAAGAAAACCGAGGTGCAGACCGGCGGCTTCAGCAAGGAGTCCATC<br>CTCCCCAAGCGCAACTCGGACAAGCTGATCGCCCGGAAGAAGGACTGG<br>GACCCGAAGAAGTACGGCGGCTTCGACAGCCCCACCGTCGCCTACTCC<br>GTGCTGGTCGTGGCGAAGGTCGAGAAGGGCAAGAGCAAGAAGCTGAAG<br>TCCGTGAAGGAGCTGCTCGGCATCACCATCATGGAGCGCTCCTCGTTCG<br>AGAAGAACCCGATCGACTTCCTGGAGGCCAAGGGCTACAAGGAGGTCA<br>AGAAGGACCTCATCATCAAGCTGCCCAAGTACTCGCTGTTCGAGCTCGA<br>GAACGGCCGCAAGCGGATGCTCGCCAGCGCGGGCGAGCTGCAGAAGG<br>GCAACGAGCTGGCCCTCCCGTCCAAGTACGTCAACTTCCTGTACCTCGC<br>GTCCCACTACGAGAAGCTGAAGGGCTCGCCCGAGGACAACGAGCAGAA<br>GCAGCTCTTCGTGGAGCAGCACAAGCACTACCTGGACGAGATCATCGAG<br>CAGATCTCGGAGTTCAGCAAGCGGGTCATCCTGGCCGACGCGAACCTC<br>GACAAGGTGCTGTCCGCCTACAACAAGCACCGCGACAAGCCGATCCGG<br>GAGCAGGCGGAGAACATCATCCACCTGTTCACCCTCACCAACCTGGGCG<br>CCCCCGCCGCGTTCAAGTACTTCGACACCACCATCGACCGCAAGCGGTA<br>CACCAGCACCAAGGAGGTCCTCGACGCGACCCTGATCCACCAGTCCATC<br>ACCGGCCTGTACGAAACCCGCATCGACCTCTCCCAGCTCGGCGGCGAC |

TGAGAATTCAGATCTACGCGTTCCCCGCAAAAGCGGCCTTTGACTCCCT
GCAAGCCTCAGCGACCGAATATATCGGTTATGCGTGGGCGATGGTTGTT
GTCATTGTCGGCGCAACTATCGGTATCAAGCTGTTTAAGAAATTCACCTC
GAAAGCAAGCTGATAAACCGATACAATTAAAGGCTCCTTTTGGAGCCTTT
TTTTGACAGCGTGCAGGACTGGGGGAGTTATGTTCACATTCGAACCGTC
TCTGCTTTGACACGGACAAGCGCTATGGTGTAAAGTCGTGGCCACATGT
TGTCCTCCTCGCCCTGTTTTAGAGCTAGAAATAGCAAGTTAAAATAAGGC
TAGTCCGTTATCAACTTGAAAAAGTGGCACCGAGTCGGTGCTTTTTACTC
CATCTGGATTTGTTCAGAACGCTCGGTTGCCGCCGGGCGTTTTTTATCTA
GAGGCCAGGAACCGTAAAAAGGCCGCGTTGCTGGCGTTTTTCCATAGGC
TCCGCCCCCCTGACGAGCATCACAAAAATCGACGCTCAAGTCAGAGGTG
GCGAAACCCGACAGGACTATAAAGATACCAGGCGTTTCCCCCTGGAAGC
TCCCTCGTGCGCTCTCCTGTTCCGACCCTGCCGCTTACCGGATACCTGT
CCGCCTTTCTCCCTTCGGGAAGCGTGGCGCTTTCTCATAGCTCACGCTG
TAGGTATCTCAGTTCGGTGTAGGTCGTTCGCTCCAAGCTGGGCTGTGTG
CACGACCCCCCGTTCAGCCCGACCGCTGCGCCTTATCCGGTAACTATC
GTCTTGAGTCCAACCCGGTAAGACACGACTTATCGCCACTGGCAGCAGC
CACTGGTAACAGGATTAGCAGAGCGAGGTATGTAGGCGGTGCTACAGAG
TTCTTGAAGTGGTGGCCTAACTACGGCTACACTAGAAGAACAGTATTTGG
TATCTGCGCTCTGCTGAAGCCAGTTACCTTCGGAAAAAGAGTTGGTAGCT
CTTGATCCGGCAAACAAACCACCGCTGGTAGCGGTGGTTTTTTTGTTTGC
AAGCAGCAGATTACGCGCAGAAAAAAAGGATCTCAAGAAGATCCTTTGAT
CTTTTCTACGGGGTCTGACGCTCAGTGGAACGAAAACTCACGTTAAGGG
ATTTTGGTCATGAGATTATCAAAAAGGATCTTCACCTAGATCCTTTTGGTT
CATGTGCAGCTCCATCAGCAAAAGGGGATGATAAGTTTATCACCACCGA
CTATTTGCAACAGTGCCGTTGATCGTGCTATGATCGACTGATGTCATCAG
CGGTGGAGTGCAATGTCGTGCAATACGAATGGCGAAAAGCCGAGCTCAT
CGGTCAGCTTCTCAACCTTGGGGTTACCCCCGGCGGTGTGCTGCTGGTC
CACAGCTCCTTCCGTAGCGTCCGGCCCCTCGAAGATGGGCCACTTGGA
CTGATCGAGGCCCTGCGTGCTGCGCTGGGTCCGGGAGGGACGCTCGTC
ATGCCCTCGTGGTCAGGTCTGGACGACGAGCCGTTCGATCCTGCCACGT
CGCCCGTTACACCGGACCTTGGAGTTGTCTCTGACACATTCTGGCGCCT
GCCAAATGTAAAGCGCAGCGCCCATCCATTTGCCTTTGCGGCAGCGGG
GCCACAGGCAGAGCAGATCATCTCTGATCCATTGCCCCTGCCACCTCAC
TCGCCTGCAAGCCCGGTCGCCCGTGTCCATGAACTCGATGGGCAGGTA
CTTCTCCTCGGCGTGGGACACGATGCCAACACGACGCTGCATCTTGCCG
AGTTGATGGCAAAGGTTCCCTATGGGGTGCCGAGACACTGCACCATTCT
TCAGGATGGCAAGTTGGTACGCGTCGATTATCTCGAGAATGACCACTGC
TGTGAGCGCTTTGCCTTGGCGGACAGGTGGCTCAAGGAGAAGAGCCTT
CAGAAGGAAGGTCCAGTCGGTCATGCCTTTGCTCGGTTGATCCGCTCCC
GCGACATTGTGGCGACAGCCCTGGGTCAACTGGGCCGAGATCCGTTGA
TCTTCCTGCATCCGCCAGAGGCGGGATGCGAAGAATGCGATGCCGCTC
GCCAGTCGATTGGCTGAGCTCATGAGCGGAGAACGAGATGACGTTGGA
GGGGCAAGGTCGCGCTGATTGCTGGGGCAACACGTGGAGCGGATCGG
GGATTGTCTTTCTTCAGCTCGCTGATGATATGCTGACGCTCAATGCCGTT
TGGCCTCCGACTAACGAAAATCCCGCATTTGGACGGCTGATCCGATTGG
CACGGCGGACGGCGAATGGCGGAGCAGACGCTCGTCCGGGGGCAATG
AGATATGAAAAAGCCTGAACTCACCGCGACGTATCGATGTCGAGGTTCC
TCAGGGGAGCCACCCCAGAGAAGCCCTCGGAGCTGAGCGGAGCTATTT
CCAAAGCCATGCCAGCTAGAGACAGTGCACACCGCCAAGCATCTGCAAA
ACCCTCGCCTGGAGGGAAAGTGCAATGTACGTACGCTGGTTTCCCTCCA

| | |
|---|---|
| | GAAAGGGATTCTGCGGCTTCTCAACTTTGGAAGAAGAGGCGGAACGAAC<br>TGCTGTTTCAGCCTACTCATGTGAGGAGGCTGGTCCTTTACCCTGATGC<br>CCGAGGGCATCAGGGTAAAGGACCAGCCTCGCCAACTAGGAGCGCTAC<br>CAAGGCGAACACAACCCAGGACTTGGTAGAACCCTTCGCCGGAGAAGTT<br>CAGACTCATCGGCATGAGCACCCCTCGCCGCGCACTCGGCACCGGTCC<br>GTCGACCACCACCAGGACGTCGTTGTCGACGTCGGCCCCGCGGCTCCT<br>GCCCGCCGAACGCGTCGTCGTCGACGGCCTGGTGCTCATCGACGAGCA<br>CCCGGAGCCAGGTGAAAAGCGCCGGCGGACGCTCGGACTGGGCGCGG<br>GATTCCAGCAGTAACCCAGGTCCGCCGGCCACCTCACGGCAGGCAGAC<br>CCTCGGCTTTCGCGCCGGGACCGCCATGAGACCGCCACCCGGATGTCC<br>GGGGTGGCGGTCTCATGGCGGTCCCTCAGCGGCCCTACGCGACCGCTG<br>TGTCGACGCGGAGGCAGTCTCCGGGGTGCTGGCCCCAGGGCTGGAGTA<br>CCGGGGCGAGCTTGCCCTTGCAGCGGCGGCACACGGGCGACGCGGCG<br>CCGGGCGGGGGAGTCTTGACGTCGACGGTGACCGGCTTCGGCTTCAGG<br>CGCTTCCGAAGGCTGATCGTCGGGCGGATCGCCTCCTTCTTCGGCTGAC<br>GCACTCGGTCGAGCGAAGCGGCCAGCTCCTCCGTGCGGGCCTCGTTGG<br>CCTTCCGGCGCGCCTCGCGGAAAGCAGCTTCCTCCTCGGACATGACCT<br>CGAACCTCATCTGGTCAGCGTCAAGGTCGCCCGGCGCGGCCGGCGCTT<br>CCGGGGCGGGCGGGTCCAGGACGTCCTTGCCCCACACCAGGCCCCAG<br>GACTCGACGAGCCGCCGGACGCCCGGTAGGCCGTACGTCTCGGCGAC<br>CTTGATGAGGTCGAGGCGACGTCCGGCGACGCGGGCGATGTATCGGTA<br>CCAGATGTAGGCCGGGATGACCGCGATGGCGACCAGGCCCTCGGTGTC<br>GTCGGTGATCTCCTCCTCGGTGCGGACGTCCTGCTGGATGCCGAGTTCC<br>TTGATCAGCCGGTTCAGGTTCTGCGACCGGTAGTGCTTGCGGACCTGGA<br>ACACGCCGAACTCGCGCTCGCGGTACTTCTCGACGAACGGGCCGGGCC<br>GACGAAGCCGCTGCAGCTCGGCGGCCGCCGCGTCGCCCAGGTCGAGC<br>GGTCCCATGCGGTCGTCGCCGCGACCGGCCTTGAAGTTCTGTCCGGCC<br>AGCTCCAGGCCGATCTTGGCGACGCCGCCCTTGGTCTTGTCGCCGTCCT<br>TGTAGAGGTAGCGGGCCTGCTTGCCCGCATCGCCGTCAGCGGCGTCCG<br>CGCCGTTGAGTGGGCGCACGTCGGTGCCGTGGCCCTTGCCCTCACAGG<br>AGCAACCGGGCCGGTCGCACGTCTCGCTGACGGTGTAGCCGCCCGCG<br>GATTCGACCCCGGCGGCCCAGGCTCCGGCGAGTGCGTCGCGGAACGC<br>GGCCTGGGCGTCCGGGCCGAGCACCTCGCGGGTGACCCAGAGCGTGT<br>GCCAGTGCAGGTGCCAGCCGGAGCCCCAGCCGAAGGTGTCCTCGAAGG<br>CCCGCTCGTAGCCGATGATCCCGAAGTCGTCGCGCATCGTGCGCCAGC<br>GGCGGCCGGACGAGCCGTACGCGCCCTTCCAGCCGTCGTGCAAGACC<br>GCGACCAGGCCGTGCCGCATTCCCTTGCGGACGGTGCCGAACGCCATG<br>CGCTCGAAGTGGCGCAACGTGTTCGTGCCAAGGTGCAGCCCGTACCCG<br>GCGTCCGCGAGACCGTCGGCGGCGAGCTGCACGTTCGAGCCCCGTAC<br>GGCCAGGATGCGGCTCATGCACCACGGGCAGGTGTGGACGTTGTTGCA<br>GCGGCACGTGTTGCCCCACGTCGCCTCGCCCGGCTTCCACATCAGCTC<br>GGCCGTCCCGGCAGTGAGCCGGGTCCCGCAGCCCTTGAACGCCTCGTT<br>CAGCGACACCGTCTGGTGCCGGTCCCGCCGGGCGAACCGCTCGTCGC<br>GCGGGTCCTCCCGCCCGGCTGTCGCGGCACCCTCGTTTGGGGTAGAAC<br>CCGTTCCAGTTACAGCGCTCTGACCTGCAGTGGACGGAGATTTTCCCTT<br>ACTACTAAAGCCCGCGTCCGGATTACCCGCTGTAGTCGTGCTTGCTACG<br>CTGCGTGACTGGTCCGCAATGAGACGCTTTGCGCGCTTTCGGCAGGCG<br>TCCGAGCAGTAGATTTTGGGGCGCTTCCCGGGGATGTGGACGATCGGG<br>GTGCCGCAGTGGCACTTCGGTCCGGCGGGGCGCGGTGGTGTCGACGC<br>GCTGTTCTCTCGTACGCTCGTCACAGAGCAAACGTCCTCACTCGGCATG<br>CTGCGCCGGTTCGGGGGCGGCGAGCCCGGGAGGCCAATCCCGGGCTC |

| | |
|---|---|
| | GTGCCATTTCTGGGTCCTGTTGATCCTGGCATTGGTGTGGCCGTTCATT<br>GCCCCTGCTCGCTCCTGACGCGCCGATAGACGTCCGATACGCCCGGTG<br>CTGGTGGGATTTGATAGGTCGGAAGAAGCCCCGCCCGGGCCTGGGCGG<br>GGCTTCCTGTGCGTCAGGACCTCCTCGTCGTGAGCCTCTTCGGCCTATG<br>GACGGAGTGACCTCGTGATCCGTTACAGCCGCGCGCGCTCGCGTAGAG<br>CGGTCTCATCAGTTCCACGAACGGTCCTCTTCGCAGATCAGGGCGTTGG<br>GGCGGAGTCTCACCAAGGACTACGTCTGCTGGCGATTTCCGTTACACCC<br>CGGGCGGTGGCCGGCGCACACGCGCGCCCGCGTTGGGCAGTGCAGAA<br>AGTGCAGAAACCTAGGCGCTGATGGTCCAGGTCCACGGTTCGTCGTCG<br>GCGGCGGCGCGGGCGGCGGCGTCGGCCAGGGCGCGGGCGAGACCGG<br>CTACGGCGGGCTTGATGCGCCGGTTGCGGGCGACCTTGAGCAGCTAGT<br>ATGCAGGTCGACGGATCTTTTCCGCTGCATAACCCTGCTTCGGGGTCAT<br>TATAGCGATTTTTTCGGTATATCCATCCTTTTTCGCACGATATACAGGATT<br>TTGCCAAAGGGTTCGTGTAGACTTTCCTTGGTGTATCCAACGGCGTCAG<br>CCGGGCAGGATAGGTGAAGTAGGCCCACCCGCGAGCGGGTGTTCCTTC<br>TTCACTGTCCCTTATTCGCACCTGGCGGTGCTCAACGGGAATCCTGCTC<br>TGCGAGGCTGGCCGGCTACCGCCGGCGTAACAGATGAGGGCAAGCGG<br>ATGGCTGATGAAACCAAGCCAACCAGGAAGGGCAGCCCACCTATCAAGG<br>TGTACTGCCTTCCAGACGAACGAAGAGCGATTGAGGAAAAGGCGGCGG<br>CGGCCGGCATGAGCCTGTCGGCCTACCTGCTGGCCGTCGGCCAGGGCT<br>ACAAAATCACGGGCGTCGTGGACTATGAGCACGTCCGCGAGCTGGACA<br>GCGTGCAGGACTGGGGGAGTTA |
| **CRISPRa**<br><br>(Promoter-RiboJ-RBS-dCas9-XTEN linker - AD -terminator-promoter-sgRNA targeting sequence-sgRNA scaffold-terminator-ori-AprR-pSG5 rep-pSG5 ds-oriT) | TGTTCACATTCGAACCGTCTCTGCTTTGACATCGTGTGGCGCTTGGGTG<br>TAAAGTCGTGGCCATTAAACAAAATTATTTGTAGAGGCTGTTTCGTCCTC<br>ACGGACTCATCAGACCGGAAAGCACATCCGGTGACAGCTAACTACGAA<br>GGGGAGTCAGTATGGACAAGAAGTACAGCATCGGCCTGGCCATCGGCA<br>CCAACAGCGTGGGCTGGGCGGTCATCACCGACGAGTACAAGGTCCCCT<br>CCAAGAAGTTCAAGGTCCTGGGCAACACCGACCGGCACTCGATCAAGA<br>AGAACCTGATCGGCGCCCTGCTCTTCGACAGCGGCGAAACCGCCGAG<br>GCGACCCGCCTGAAGCGGACCGCCCGTCGCCGCTACACCCGGCGCAA<br>GAACCGCATCTGCTACCTGCAGGAGATCTTCTCCAACGAGATGGCCAA<br>GGTCGACGACTCGTTCTTCCACCGGCTCGAGGAGAGCTTCCTGGTGGA<br>GGAGGACAAGAAGCACGAGCGCCACCCGATCTTCGGCAACATCGTCGA<br>CGAGGTGGCCTACCACGAGAAGTACCCCACCATCTACCACCTCCGCAA<br>GAAGCTGGTCGACTCGACCGACAAGGCGGACCTGCGGCTCATCTACCT<br>GGCCCTCGCGCACATGATCAAGTTCCGCGGCCACTTCCTCATCGAGGG<br>CGACCTGAACCCGGACAACTCCGACGTCGACAAGCTCTTCATCCAGCT<br>GGTGCAGACCTACAACCAGCTGTTCGAGGAGAACCCCATCAACGCCAG<br>CGGCGTCGACGCCAAGGCGATCCTCTCCGCGCGCCTGAGCAAGTCCC<br>GGCGCCTGGAGAACCTCATCGCCCAGCTGCCGGGCGAGAAGAAGAAC<br>GGCCTCTTCGGCAACCTGATCGCGCTGTCGCTCGGCCTGACCCCCAAC<br>TTCAAGAGCAACTTCGACCTGGCCGAGGACGCGAAGCTCCAGCTGTCC<br>AAGGACACCTACGACGACGACCTGGACAACCTGCTCGCCCAGATCGGC<br>GACCAGTACGCGGACCTCTTCCTGGCCGCGAAGAACCTCTCGGACGCC<br>ATCCTGCTCAGCGACATCCTGCGGGTCAACACCGAGATCACCAAGGCC<br>CCGCTGTCGGCGAGCATGATCAAGCGGTACGACGAGCACCACCAGGA<br>CCTGACCCTGCTCAAGGCCCTCGTGCGCCAGCAGCTGCCCGAGAAGTA<br>CAAGGAGATCTTCTTCGACCAGTCCAAGAACGGCTACGCCGGCTACAT<br>CGACGGCGGCGCGTCGCAGGAGGAGTTCTACAAGTTCATCAAGCCGAT<br>CCTGGAGAAGATGGACGGCACCGAGGAGCTGCTCGTCAAGCTGAACC<br>GCGAGGACCTGCTCCGCAAGCAGCGGACCTTCGACAACGGCTCCATCC |

| | |
|---|---|
| | CGCACCAGATCCACCTGGGCGAGCTCCACGCCATCCTCCGGCGCCAG<br>GAGGACTTCTACCCCTTCCTGAAGGACAACCGCGAGAAGATCGAGAAG<br>ATCCTGACCTTCCGGATCCCGTACTACGTCGGCCCCCTGGCCCGCGGC<br>AACTCCCGGTTCGCGTGGATGACCCGGAAGTCGGAGGAAACCATCACC<br>CCGTGGAACTTCGAGGAGGTCGTGGACAAGGGCGCCTCCGCGCAGTC<br>GTTCATCGAGCGCATGACCAACTTCGACAAGAACCTCCCGAACGAGAA<br>GGTCCTGCCCAAGCACAGCCTGCTCTACGAGTACTTCACCGTGTACAA<br>CGAGCTGACCAAGGTCAAGTACGTGACCGAGGGCATGCGGAAGCCGG<br>CCTTCCTGTCCGGCGAGCAGAAGAAGGCGATCGTCGACCTGCTCTTCA<br>AGACCAACCGCAAGGTCACCGTGAAGCAGCTGAAGGAGGACTACTTCA<br>AGAAGATCGAGTGCTTCGACTCCGTCGAGATCTCGGGCGTGGAGGACC<br>GCTTCAACGCCTCCCTGGGCACCTACCACGACCTGCTCAAGATCATCAA<br>GGACAAGGACTTCCTCGACAACGAGGAGAACGAGGACATCCTGGAGGA<br>CATCGTCCTCACCCTGACCCTCTTCGAGGACCGCGAGATGATCGAGGA<br>GCGGCTCAAGACCTACGCCCACCTGTTCGACGACAAGGTGATGAAGCA<br>GCTGAAGCGGCGCCGGTACACCGGCTGGGGCCGCCTCTCCCGGAAGC<br>TGATCAACGGCATCCGGGACAAGCAGAGCGGCAAGACCATCCTGGACT<br>TCCTCAAGTCCGACGGCTTCGCCAACCGCAACTTCATGCAGCTCATCCA<br>CGACGACTCGCTGACCTTCAAGGAGGACATCCAGAAGGCCCAGGTGTC<br>CGGCCAGGGCGACAGCCTCCACGAGCACATCGCCAACCTGGCGGGCT<br>CCCCGGCGATCAAGAAGGGCATCCTCCAGACCGTCAAGGTCGTGGAC<br>GAGCTGGTCAAGGTGATGGGCCGCCACAAGCCCGAGAACATCGTGATC<br>GAGATGGCCCGGGAGAACCAGACCACCCAGAAGGGCCAGAAGAACTC<br>CCGCGAGCGGATGAAGCGCATCGAGGAGGGCATCAAGGAGCTCGGCT<br>CGCAGATCCTGAAGGAGCACCCGGTCGAGAACACCCAGCTCCAGAACG<br>AGAAGCTGTACCTCTACTACCTGCAGAACGGCCGCGACATGTACGTGG<br>ACCAGGAGCTCGACATCAACCGGCTGAGCGACTACGACGTCGACGCCA<br>TCGTGCCGCAGTCCTTCCTGAAGGACGACTCGATCGACAACAAGGTCC<br>TGACCCGCTCCGACAAGAACCGGGGCAAGTCCGACAACGTGCCCTCG<br>GAGGAGGTCGTGAAGAAGATGAAGAACTACTGGCGCCAGCTGCTCAAC<br>GCCAAGCTCATCACCCAGCGCAAGTTCGACAACCTGACCAAGGCCGAG<br>CGGGGCGGCCTGTCGGAGCTCGACAAGGCGGGCTTCATCAAGCGCCA<br>GCTCGTCGAAACCCGGCAGATCACCAAGCACGTGGCCCAGATCCTGGA<br>CAGCCGGATGAACACCAAGTACGACGAGAACGACAAGCTGATCCGCGA<br>GGTCAAGGTGATCACCCTCAAGAGCAAGCTGGTGTCCGACTTCCGCAA<br>GGACTTCCAGTTCTACAAGGTCCGGGAGATCAACAACTACCACCACGC<br>CCACGACGCGTACCTGAACGCCGTCGTGGGCACCGCGCTGATCAAGAA<br>GTACCCGAAGCTGGAGTCCGAGTTCGTCTACGGCGACTACAAGGTCTA<br>CGACGTGCGCAAGATGATCGCCAAGTCGGAGCAGGAGATCGGCAAGG<br>CCACCGCGAAGTACTTCTTCTACAGCAACATCATGAACTTCTTCAAGAC<br>CGAGATCACCCTGGCCAACGGCGAGATCCGCAAGCGGCCCCTGATCG<br>AAACCAACGGCGAAACCGGCGAGATCGTCTGGGACAAGGGCCGCGAC<br>TTCGCCACCGTCCGGAAGGTGCTGTCCATGCCGCAGGTCAACATCGTC<br>AAGAAAACCGAGGTGCAGACCGGCGGCTTCAGCAAGGAGTCCATCCTC<br>CCCAAGCGCAACTCGGACAAGCTGATCGCCCGGAAGAAGGACTGGGA<br>CCCGAAGAAGTACGGCGGCTTCGACAGCCCCACCGTCGCCTACTCCGT<br>GCTGGTCGTGGCGAAGGTCGAGAAGGGCAAGAGCAAGAAGCTGAAGT<br>CCGTGAAGGAGCTGCTCGGCATCACCATCATGGAGCGCTCCTCGTTCG<br>AGAAGAACCCGATCGACTTCCTGGAGGCCAAGGGCTACAAGGAGGTCA<br>AGAAGGACCTCATCATCAAGCTGCCCAAGTACTCGCTGTTCGAGCTCGA<br>GAACGGCCGCAAGCGGATGCTCGCCAGCGCGGGCGAGCTGCAGAAGG |

| | |
|---|---|
| | GCAACGAGCTGGCCCTCCCGTCCAAGTACGTCAACTTCCTGTACCTCG<br>CGTCCCACTACGAGAAGCTGAAGGGCTCGCCCGAGGACAACGAGCAG<br>AAGCAGCTCTTCGTGGAGCAGCACAAGCACTACCTGGACGAGATCATC<br>GAGCAGATCTCGGAGTTCAGCAAGCGGGTCATCCTGGCCGACGCGAAC<br>CTCGACAAGGTGCTGTCCGCCTACAACAAGCACCGCGACAAGCCGATC<br>CGGGAGCAGGCGGAGAACATCATCCACCTGTTCACCCTCACCAACCTG<br>GGCGCCCCCGCCGCGTTCAAGTACTTCGACACCACCATCGACCGCAAG<br>CGGTACACCAGCACCAAGGAGGTCCTCGACGCGACCCTGATCCACCAG<br>TCCATCACCGGCCTGTACGAAACCCGCATCGACCTCTCCCAGCTCGGC<br>GGCGACTGAAGTGGCTCAGAGACGCCGGGTACTTCTGAGTCCGCTACG<br>CCTGAGAGCATGCTTATCGCTCAGCGTCCTTCGCTGACCGAAGAGGTC<br>GTCGACGAGTTCCGCTCCCGGTTCGTGATCGAGCCGCTGGAGCCGGG<br>CTTCGGCTACACCCTCGGCAACTCCCTCCGCCGTACCCTCCTCTCCTC<br>GATCCCGGGTGCCGCTGTCACCAGCATCCGCATCGACGGTGTCCTGCA<br>CGAGTTCACCACCGTGCCGGGCGTCAAGGAGGACGTCACCGACCTCAT<br>CCTCAACATCAAGCAGCTGGTCGTCTCCTCGGAGCACGACGAGCCGGT<br>CGTGATGTACCTGCGCAAGCAGGGCCCGGGTCTGGTCACCGCCGCCG<br>ACATCGCGCCCCCGGCCGGTGTCGAGGTGCACAACCCCGACCTCGTC<br>CTCGCCACGCTCAACGGCAAGGGCAAGCTGGAGATGGAGCTGACCGT<br>CGAGCGCGGTCGCGGCTACGTCTCCGCCGTGCAGAACAAGCAGGTCG<br>GTCAGGAGATCGGGCGCATCCCGGTCGACTCGATCTACTCGCCGGTTC<br>TCAAGGTCACCTACAAGGTCGAGGCGACCCGAGTCGAGCAGCGCACC<br>GACTTCGACAAGCTGATCGTCGACGTCGAGACCAAGCAGGCCATGCGC<br>CCGCGTGACGCCATGGCGTCCGCCGGCAAGACCCTGGTCGAGCTGTT<br>CGGTCTGGCGCGCGAGCTCAACATCGACGCCGAATTCAGATCTACGCG<br>TTCCCCGCAAAAGCGGCCTTTGACTCCCTGCAAGCCTCAGCGACCGAA<br>TATATCGGTTATGCGTGGGCGATGGTTGTTGTCATTGTCGGCGCAACTA<br>TCGGTATCAAGCTGTTTAAGAAATTCACCTCGAAAGCAAGCTGATAAAC<br>CGATACAATTAAAGGCTCCTTTTGGAGCCTTTTTTTGACAGCGTGCAGG<br>ACTGGGGGAGTTATGTTCACATTCGAACCGTCTCTGCTTTGACACGGAC<br>AAGCGCTATGGTGTAAAGTCGTGGCCACGTCAGAATTCACAAGCCCGG<br>TTTTAGAGCTAGAAATAGCAAGTTAAAATAAGGCTAGTCCGTTATCAACT<br>TGAAAAAGTGGCACCGAGTCGGTGCTTTTTACTCCATCTGGATTTGTTC<br>AGAACGCTCGGTTGCCGCCGGGCGTTTTTTATCTAGAGGCCAGGAACC<br>GTAAAAAGGCCGCGTTGCTGGCGTTTTTCCATAGGCTCCGCCCCCCTG<br>ACGAGCATCACAAAAATCGACGCTCAAGTCAGAGGTGGCGAAACCCGA<br>CAGGACTATAAAGATACCAGGCGTTTCCCCCTGGAAGCTCCCTCGTGC<br>GCTCTCCTGTTCCGACCCTGCCGCTTACCGGATACCTGTCCGCCTTTCT<br>CCCTTCGGGAAGCGTGGCGCTTTCTCATAGCTCACGCTGTAGGTATCTC<br>AGTTCGGTGTAGGTCGTTCGCTCCAAGCTGGGCTGTGTGCACGACCCC<br>CCCGTTCAGCCCGACCGCTGCGCCTTATCCGGTAACTATCGTCTTGAGT<br>CCAACCCGGTAAGACACGACTTATCGCCACTGGCAGCAGCCACTGGTA<br>ACAGGATTAGCAGAGCGAGGTATGTAGGCGGTGCTACAGAGTTCTTGA<br>AGTGGTGGCCTAACTACGGCTACACTAGAAGAACAGTATTTGGTATCTG<br>CGCTCTGCTGAAGCCAGTTACCTTCGGAAAAAGAGTTGGTAGCTCTTGA<br>TCCGGCAAACAAACCACCGCTGGTAGCGGTGGTTTTTTTGTTTGCAAGC<br>AGCAGATTACGCGCAGAAAAAAAGGATCTCAAGAAGATCCTTTGATCTT<br>TTCTACGGGGTCTGACGCTCAGTGGAACGAAAACTCACGTTAAGGGATT<br>TTGGTCATGAGATTATCAAAAAGGATCTTCACCTAGATCCTTTTGGTTCA<br>TGTGCAGCTCCATCAGCAAAAGGGGATGATAAGTTTATCACCACCGACT<br>ATTTGCAACAGTGCCGTTGATCGTGCTATGATCGACTGATGTCATCAGC |

| | |
|---|---|
| | GGTGGAGTGCAATGTCGTGCAATACGAATGGCGAAAAGCCGAGCTCAT<br>CGGTCAGCTTCTCAACCTTGGGGTTACCCCCGGCGGTGTGCTGCTGGT<br>CCACAGCTCCTTCCGTAGCGTCCGGCCCCTCGAAGATGGGCCACTTGG<br>ACTGATCGAGGCCCTGCGTGCTGCGCTGGGTCCGGGAGGGACGCTCG<br>TCATGCCCTCGTGGTCAGGTCTGGACGACGAGCCGTTCGATCCTGCCA<br>CGTCGCCCGTTACACCGGACCTTGGAGTTGTCTCTGACACATTCTGGC<br>GCCTGCCAAATGTAAAGCGCAGCGCCCATCCATTTGCCTTTGCGGCAG<br>CGGGGCCACAGGCAGAGCAGATCATCTCTGATCCATTGCCCCTGCCAC<br>CTCACTCGCCTGCAAGCCCGGTCGCCCGTGTCCATGAACTCGATGGGC<br>AGGTACTTCTCCTCGGCGTGGGACACGATGCCAACACGACGCTGCATC<br>TTGCCGAGTTGATGGCAAAGGTTCCCTATGGGGTGCCGAGACACTGCA<br>CCATTCTTCAGGATGGCAAGTTGGTACGCGTCGATTATCTCGAGAATGA<br>CCACTGCTGTGAGCGCTTTGCCTTGGCGGACAGGTGGCTCAAGGAGAA<br>GAGCCTTCAGAAGGAAGGTCCAGTCGGTCATGCCTTTGCTCGGTTGAT<br>CCGCTCCCGCGACATTGTGGCGACAGCCCTGGGTCAACTGGGCCGAG<br>ATCCGTTGATCTTCCTGCATCCGCCAGAGGCGGGATGCGAAGAATGCG<br>ATGCCGCTCGCCAGTCGATTGGCTGAGCTCATGAGCGGAGAACGAGAT<br>GACGTTGGAGGGGCAAGGTCGCGCTGATTGCTGGGGCAACACGTGGA<br>GCGGATCGGGGATTGTCTTTCTTCAGCTCGCTGATGATATGCTGACGCT<br>CAATGCCGTTTGGCCTCCGACTAACGAAAATCCCGCATTTGGACGGCT<br>GATCCGATTGGCACGGCGGACGGCGAATGGCGGAGCAGACGCTCGTC<br>CGGGGGCAATGAGATATGAAAAAGCCTGAACTCACCGCGACGTATCGA<br>TGTCGAGGTTCCTCAGGGGAGCCACCCCAGAGAAGCCCTCGGAGCTG<br>AGCGGAGCTATTTCCAAAGCCATGCCAGCTAGAGACAGTGCACACCGC<br>CAAGCATCTGCAAAACCCTCGCCTGGAGGGAAAGTGCAATGTACGTAC<br>GCTGGTTTCCCTCCAGAAAGGGATTCTGCGGCTTCTCAACTTTGGAAGA<br>AGAGGCGGAACGAACTGCTGTTTCAGCCTACTCATGTGAGGAGGCTGG<br>TCCTTTACCCTGATGCCCGAGGGCATCAGGGTAAAGGACCAGCCTCGC<br>CAACTAGGAGCGCTACCAAGGCGAACACAACCCAGGACTTGGTAGAAC<br>CCTTCGCCGGAGAAGTTCAGACTCATCGGCATGAGCACCCCTCGCCGC<br>GCACTCGGCACCGGTCCGTCGACCACCACCAGGACGTCGTTGTCGAC<br>GTCGGCCCCGCGGCTCCTGCCCGCCGAACGCGTCGTCGTCGACGGCC<br>TGGTGCTCATCGACGAGCACCCGGAGCCAGGTGAAAAGCGCCGGCGG<br>ACGCTCGGACTGGGCGCGGGATTCCAGCAGTAACCCAGGTCCGCCGG<br>CCACCTCACGGCAGGCAGACCCTCGGCTTTCGCGCCGGGACCGCCAT<br>GAGACCGCCACCCGGATGTCCGGGGTGGCGGTCTCATGGCGGTCCCT<br>CAGCGGCCCTACGCGACCGCTGTGTCGACGCGGAGGCAGTCTCCGGG<br>GTGCTGGCCCCAGGGCTGGAGTACCGGGGCGAGCTTGCCCTTGCAGC<br>GGCGGCACACGGGCGACGCGGCGCCGGGCGGGGGAGTCTTGACGTC<br>GACGGTGACCGGCTTCGGCTTCAGGCGCTTCCGAAGGCTGATCGTCG<br>GGCGGATCGCCTCCTTCTTCGGCTGACGCACTCGGTCGAGCGAAGCG<br>GCCAGCTCCTCCGTGCGGGCCTCGTTGGCCTTCCGGCGCGCCTCGCG<br>GAAAGCAGCTTCCTCCTCGGACATGACCTCGAACCTCATCTGGTCAGC<br>GTCAAGGTCGCCCGGCGCGGCCGGCGCTTCCGGGGCGGGCGGGTCC<br>AGGACGTCCTTGCCCCACACCAGGCCCCAGGACTCGACGAGCCGCCG<br>GACGCCCGGTAGGCCGTACGTCTCGGCGACCTTGATGAGGTCGAGGC<br>GACGTCCGGCGACGCGGGCGATGTATCGGTACCAGATGTAGGCCGGG<br>ATGACCGCGATGGCGACCAGGCCCTCGGTGTCGTCGGTGATCTCCTCC<br>TCGGTGCGGACGTCCTGCTGGATGCCGAGTTCCTTGATCAGCCGGTTC<br>AGGTTCTGCGACCGGTAGTGCTTGCGGACCTGGAACACGCCGAACTCG<br>CGCTCGCGGTACTTCTCGACGAACGGGCCGGGCCGACGAAGCCGCTG |

| | |
|---|---|
| | CAGCTCGGCGGCCGCCGCGTCGCCCAGGTCGAGCGGTCCCATGCGGT<br>CGTCGCCGCGACCGGCCTTGAAGTTCTGTCCGGCCAGCTCCAGGCCG<br>ATCTTGGCGACGCCGCCCTTGGTCTTGTCGCCGTCCTTGTAGAGGTAG<br>CGGGCCTGCTTGCCCGCATCGCCGTCAGCGGCGTCCGCGCCGTTGAG<br>TGGGCGCACGTCGGTGCCGTGGCCCTTGCCCTCACAGGAGCAACCGG<br>GCCGGTCGCACGTCTCGCTGACGGTGTAGCCGCCCGCGGATTCGACC<br>CCGGCGGCCCAGGCTCCGGCGAGTGCGTCGCGGAACGCGGCCTGGG<br>CGTCCGGGCCGAGCACCTCGCGGGTGACCCAGAGCGTGTGCCAGTGC<br>AGGTGCCAGCCGGAGCCCCAGCCGAAGGTGTCCTCGAAGGCCCGCTC<br>GTAGCCGATGATCCCGAAGTCGTCGCGCATCGTGCGCCAGCGGCGGC<br>CGGACGAGCCGTACGCGCCCTTCCAGCCGTCGTGCAAGACCGCGACC<br>AGGCCGTGCCGCATTCCCTTGCGGACGGTGCCGAACGCCATGCGCTC<br>GAAGTGGCGCAACGTGTTCGTGCCAAGGTGCAGCCCGTACCCGGCGT<br>CCGCGAGACCGTCGGCGGCGAGCTGCACGTTCGAGCCCCGTACGGCC<br>AGGATGCGGCTCATGCACCACGGGCAGGTGTGGACGTTGTTGCAGCG<br>GCACGTGTTGCCCCACGTCGCCTCGCCCGGCTTCCACATCAGCTCGGC<br>CGTCCCGGCAGTGAGCCGGGTCCCGCAGCCCTTGAACGCCTCGTTCA<br>GCGACACCGTCTGGTGCCGGTCCCGCCGGGCGAACCGCTCGTCGCGC<br>GGGTCCTCCCGCCCGGCTGTCGCGGCACCCTCGTTTGGGGTAGAACC<br>CGTTCCAGTTACAGCGCTCTGACCTGCAGTGGACGGAGATTTTCCCTTA<br>CTACTAAAGCCCGCGTCCGGATTACCCGCTGTAGTCGTGCTTGCTACG<br>CTGCGTGACTGGTCCGCAATGAGACGCTTTGCGCGCTTTCGGCAGGCG<br>TCCGAGCAGTAGATTTTGGGGCGCTTCCCGGGGATGTGGACGATCGGG<br>GTGCCGCAGTGGCACTTCGGTCCGGCGGGGCGCGGTGGTGTCGACGC<br>GCTGTTCTCTCGTACGCTCGTCACAGAGCAAACGTCCTCACTCGGCATG<br>CTGCGCCGGTTCGGGGGCGGCGAGCCCGGGAGGCCAATCCCGGGCT<br>CGTGCCATTTCTGGGTCCTGTTGATCCTGGCATTGGTGTGGCCGTTCAT<br>TGCCCCTGCTCGCTCCTGACGCGCCGATAGACGTCCGATACGCCCGGT<br>GCTGGTGGGATTTGATAGGTCGGAAGAAGCCCCGCCCGGGCCTGGGC<br>GGGGCTTCCTGTGCGTCAGGACCTCCTCGTCGTGAGCCTCTTCGGCCT<br>ATGGACGGAGTGACCTCGTGATCCGTTACAGCCGCGCGCGCTCGCGTA<br>GAGCGGTCTCATCAGTTCCACGAACGGTCCTCTTCGCAGATCAGGGCG<br>TTGGGGCGGAGTCTCACCAAGGACTACGTCTGCTGGCGATTTCCGTTA<br>CACCCCGGGCGGTGGCCGGCGCACACGCGCGCCCGCGTTGGGCAGT<br>GCAGAAAGTGCAGAAACCTAGGCGCTGATGGTCCAGGTCCACGGTTCG<br>TCGTCGGCGGCGGCGCGGGCGGCGGCGTCGGCCAGGGCGCGGGCG<br>AGACCGGCTACGGCGGGCTTGATGCGCCGGTTGCGGGCGACCTTGAG<br>CAGCTAGTATGCAGGTCGACGGATCTTTTCCGCTGCATAACCCTGCTTC<br>GGGGTCATTATAGCGATTTTTTCGGTATATCCATCCTTTTTCGCACGATA<br>TACAGGATTTTGCCAAAGGGTTCGTGTAGACTTTCCTTGGTGTATCCAA<br>CGGCGTCAGCCGGGCAGGATAGGTGAAGTAGGCCCACCCGCGAGCGG<br>GTGTTCCTTCTTCACTGTCCCTTATTCGCACCTGGCGGTGCTCAACGGG<br>AATCCTGCTCTGCGAGGCTGGCCGGCTACCGCCGGCGTAACAGATGAG<br>GGCAAGCGGATGGCTGATGAAACCAAGCCAACCAGGAAGGGCAGCCC<br>ACCTATCAAGGTGTACTGCCTTCCAGACGAACGAAGAGCGATTGAGGA<br>AAAGGCGGCGGCGGCCGGCATGAGCCTGTCGGCCTACCTGCTGGCCG<br>TCGGCCAGGGCTACAAAATCACGGGCGTCGTGGACTATGAGCACGTCC<br>GCGAGCTGGACAGCGTGCAGGACTGGGGGAGTTA |

**Supplementary table 3. Promoters used in this study.** Plasmid sequences can be constructed by replacing the yellow regions in the example plasmids shown above. Promoter sequences were derived from Bai et al.[1], Myronovskyi and Luzhetskyy[2], and Phelan et al[3].

| Promoter ID | Sequence (5' to 3') |
|---|---|
| 57 | TTGAACGGCTGGAGGGATACACCTGGTCATAGGATACCATC |
| ermE*p | CTCTAGTATGCATGCGAGTGTCCGTTCGAGTGGCGGCTTGCGCCCGATGCTAG TCGCGGTTGATCGGCGATCGCAGGTGCACGCGGTCGATCTTGACGGCTGGCG AGAGGTGCGGGGAGGATCTGACCGACGCGGTCCACACGTGGCACCGCGATGC TGTTGTGGGCACAATCGTGCCGGTTGGTAGGATCCACAT |
| gapdh(EL) | GCTGCTCCTTCGGTCGGACGTGCGTCTACGGGCACCTTACCGCAGCCGTCGG CTGTGCGACACGGACGGATCGGGCGAACTGGCCGATGCTGGGAGAAGCGCGC TGCTGTACGGCGCGCACCGGGTGCGGAGCCCCTCGGCGAGCGGTGTGAAACT TCTGTGAATGGCCTGTTCGGTTGCTTTTTTATACGGCTGCCAGATAAGGCTTG CAGCATCTGGGCGGCTACCGCTATGATCGGGGCGTTCCTGCAATTCTTAGTGC GAGTATCTGAAAGGGGATACGC |
| KasO*p | TGTTCACATTCGAACGGTCTCTGCTTTGACAACATGCTGTGCGGTGTTGTAAAG TCGTGGCC |
| rpsl(XC) | GCCCTGCAGGCGGAAGTCAGGTAGACACGACTTCCGCTAGTCCTTGCAAGGTC TGCTGACGTGAGGCGGGGCGGTCGTTTTTGACCGCCCTGCCTTCGTCATGTAG GCTCGCTCGCTGTGCCTGGCGTGTCATCAGACGCCCAGGTCCCGGTGCCGTG AGGCCCGGGCCATCGAGCCGGTGGTACGTGGCTGCGGTCCCCTTGTGAGGGC TGCGCGCCGTGTGCTGTCCGGCGCGCACAGCCTTGAATCCACCCGCGGGGGC CGGCCGGTCTCCGTGAGCTCGAGTAGACGACGGAGACGTA |
| SP1 | TGTTCACATTCGAACCGTCTCTGCTTTGACATGGAGAGAAGTTTTGTAAAGTCG TGGCCA |
| SP10 | TGTTCACATTCGAACCGTCTCTGCTTTGACATGTTCTTACGGTCACATGTAAAGT CGTGGCCA |
| SP20 | TGTTCACATTCGAACCGTCTCTGCTTTGACACATGACGCTCACCCGTTGTAAAG TCGTGGCCA |
| SP30 | TGTTCACATTCGAACCGTCTCTGCTTTGACATCGTGTGGCGCTTGGGTGTAAAG TCGTGGCCA |
| SP43 | TGTTCACATTCGAACCGTCTCTGCTTTGACACGGACAAGCGCTATGGTGTAAAG TCGTGGCCA |

**Supplementary table 4. sgRNA sequences used in this study.** Plasmid sequences can be constructed by replacing the dark yellow regions in the example plasmids shown above.

| Sequence (5' to 3') | PAM location | Strand | Target | Type | Tool | Plasmid |
|---|---|---|---|---|---|---|
| CATGTTG TCCTCCT CGCCCT | 11 | NT | mCherry | On-target | CRISPRi | pJEC711, pJEC712, pJEC713, pJEC714 |
| TCTGGGT GCCCTCG TACGGC | 123 | NT | mCherry | On-target | CRISPRi | pJEC727 |
| GTCGGCC GGGTGCT TGACGT | 230 | NT | mCherry | On-target | CRISPRi | pJEC728 |

| | | | | | | |
|---|---|---|---|---|---|---|
| TCTTGAC TTCGGCG TCGTAG | 531 | NT | mCherry | On-target | CRISPRi | pJEC729 |
| CGTGTAG TCCTCGT TGTGCG | 623 | NT | mCherry | On-target | CRISPRi | pJEC730 |
| CAAGGGC GAGGAG GACAACA | 29 | T | mCherry | On-target | CRISPRi | pJEC731 |
| GGGGAG GGCCGG CCGTACG A | 132 | T | mCherry | On-target | CRISPRi | pJEC732 |
| GAAGGCC TACGTCA AGCACC | 242 | T | mCherry | On-target | CRISPRi | pJEC733 |
| CGAAGTC AAGACGA CGTACA | 560 | T | mCherry | On-target | CRISPRi | pJEC734 |
| CAACGAG GACTACA CGATCG | 647 | T | mCherry | On-target | CRISPRi | pJEC735 |
| GGCTCAG GTGAAGA GCGGGG | NA | NT | Non-coding genomic sequence | Off-target | CRISPRi, CRISPRa | pJEC723, pJEC737, pJEC740, pJEC743 |
| GCGTGC GTCGCAC CTCCGTG | NA | NT | Non-coding genomic sequence | Off-target | CRISPRi | pJEC724 |
| TGCACTG CTGTAAG GACGAT | NA | NA | No match | Off-target | CRISPRi | pJEC725 |
| CCGTGTG GCCCGCT CTTGTT | 44 | NT | jadR2 | On-target | CRISPRi | pJEC749 |
| AGGTATC CTGCGGT GTCCTG | -82 | T | mCherry | On-target | CRISPRa | pJEC739, pJEC742, pJEC745 |
| AGGACGC CTTTGGT AACCGC | -83 | NT | mCherry | On-target | CRISPRa | pJEC738, pJEC741, pJEC744 |
| ATGTGAA CAAAGGA CGCCTT | -73 | NT | mCherry | On-target | CRISPRa | pJEC747 |
| TGGTAAC CGCAGGA CACCGC | -93 | NT | mCherry | On-target | CRISPRa | pJEC748 |
| CGTCAGA ATTCACA AGCCCG | -73 | NT | jadJ | On-target | CRISPRa | pJEC750 |

**Supplementary table 5. Activator domains (ADs) used in this study.** Plasmid sequences can be constructed by replacing the dark green regions in the example plasmids shown above.

| AD | Nucleotide sequence | Protein sequence | Plasmids |
|---|---|---|---|
| αNTD | ATGCTTATCGCTCAGCGTCCTTCGCTGA CCGAAGAGGTCGTCGACGAGTTCCGCT CCCGGTTCGTGATCGAGCCGCTGGAGC CGGGCTTCGGCTACACCCTCGGCAACTC CCTCCGCCGTACCCTCCTCTCCTCGATC CCGGGTGCCGCTGTCACCAGCATCCGC ATCGACGGTGTCCTGCACGAGTTCACCA CCGTGCCGGGCGTCAAGGAGGACGTCA CCGACCTCATCCTCAACATCAAGCAGCT GGTCGTCTCCTCGGAGCACGACGAGCC GGTCGTGATGTACCTGCGCAAGCAGGG CCCGGGTCTGGTCACCGCCGCCGACAT CGCGCCCCCGGCCGGTGTCGAGGTGCA CAACCCCGACCTCGTCCTCGCCACGCTC AACGGCAAGGGCAAGCTGGAGATGGAG CTGACCGTCGAGCGCGGTCGCGGCTAC GTCTCCGCCGTGCAGAACAAGCAGGTC GGTCAGGAGATCGGGCGCATCCCGGTC GACTCGATCTACTCGCCGGTTCTCAAGG TCACCTACAAGGTCGAGGCGACCCGAGT CGAGCAGCGCACCGACTTCGACAAGCT GATCGTCGACGTCGAGACCAAGCAGGC CATGCGCCCGCGTGACGCCATGGCGTC CGCCGGCAAGACCCTGGTCGAGCTGTT CGGTCTGGCGCGCGAGCTCAACATCGA CGCC | MLIAQRPSLTEEVVDE FRSRFVIEPLEPGFGY TLGNSLRRTLLSSIPGA AVTSIRIDGVLHEFTTV PGVKEDVTDLILNIKQL VVSSEHDEPVVMYLR KQGPGLVTAADIAPPA GVEVHNPDLVLATLNG KGKLEMELTVERGRG YVSAVQNKQVGQEIG RIPVDSIYSPVLKVTYK VEATRVEQRTDFDKLI VDVETKQAMRPRDAM ASAGKTLVELFGLARE LNIDA | pJEC737, pJEC738, pJEC739, pJEC747, pJEC748, pJEC750 |
| ω | GTGTCCTCTTCCATCACCGCGCCCGAGG GCATCATCAACCCGCCGATCGACGAGCT GCTCGAGGCCACGGACTCGAAGTACAG CCTCGTGATCTACGCGGCCAAGCGCGC GCGTCAGATCAACGCGTACTACTCCCAG CTCGGCGAGGGCCTGCTGGAGTACGTG GGTCCGCTGGTCGACACCCACGTCCAC GAGAAGCCGCTCTCGATCGCCCTGCGC GAGATCAACGCGGGCCTGCTGACGTCC GAGGCCATCGAGGGCCCGGCGCAG | VSSSITAPEGIINPPIDE LLEATDSKYSLVIYAAK RARQINAYYSQLGEGL LEYVGPLVDTHVHEKP LSIALREINAGLLTSEAI EGPAQ | pJEC740, pJEC741, pJEC742 |
| RbpA | ATGAGTGAGCGAGCTCTTCGCGGCACG CGCCTCGTGGTGACGAGCTACGAGACC GACCGCGGCATCGATCTGGCCCCGCGC CAGGCCGTGGAGTACGCATGCGAGAAG GGCCATCGTTTTGAGATGCCCTTCTCGG TGGAAGCGGAAATTCCGCCGGAGTGGG AGTGCAAGGTCTGCGGAATCCAGGCACT CCTGGTGGACGGGGACGGACCTGAGGA | MSERALRGTRLVVTSY ETDRGIDLAPRQAVEY ACEKGHRFEMPFSVE AEIPPEWECKVCGIQA LLVDGDGPEEKKGKP ARTHWDMLMERRTRE ELEEVLAERLAVLRSG AMNIAVHPRDSRKSA | pJEC743, pJEC744, pJEC745 |

| | |
|---|---|
| | GAAGAAGGGCAAGCCTGCGCGTACGCA CTGGGACATGCTCATGGAGCGACGCAC CCGCGAGGAGCTGGAGGAGGTCCTCGC CGAAAGGCTGGCCGTCCTGCGTTCCGG CGCCATGAACATCGCCGTGCATCCGCG CGACAGCCGCAAGTCCGCC |

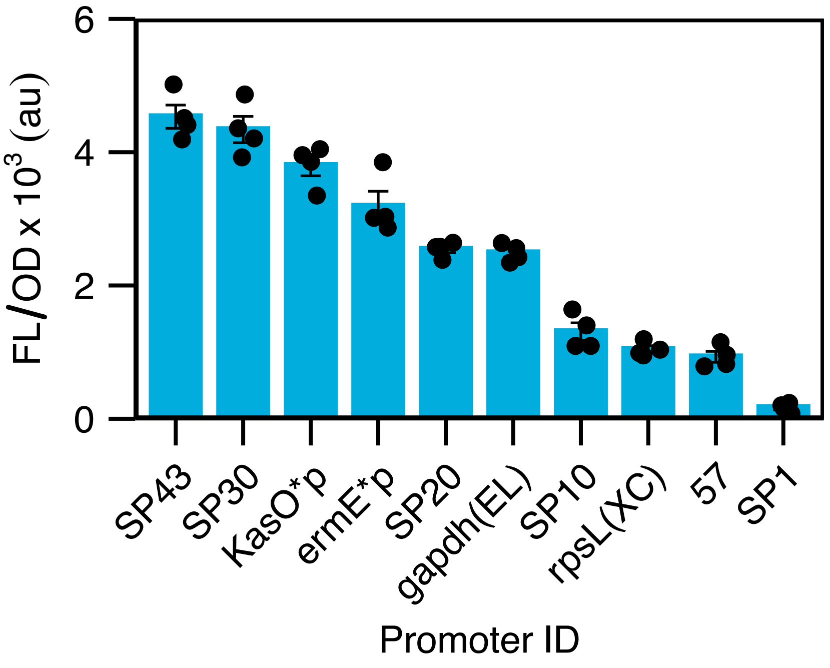

**Supplementary figure 1. Evaluating the strength of a library of *Streptomyces* promoters.** Fluorescence characterization of constitutive promoters. The expression strength of a constitutive promoter library was evaluated by cloning each promoter upstream of an mCherry reporter and integrating the resulting reporter constructs into the genome of *S. venezuelae* cells at the ΦC31 attB site. Fluorescence characterization was performed by bulk fluorescence measurements (measured in units of fluorescence [FL]/optical density [OD] at 600 nm). Data represent mean values and errors bars represent standard deviation of 4 biological replicates.

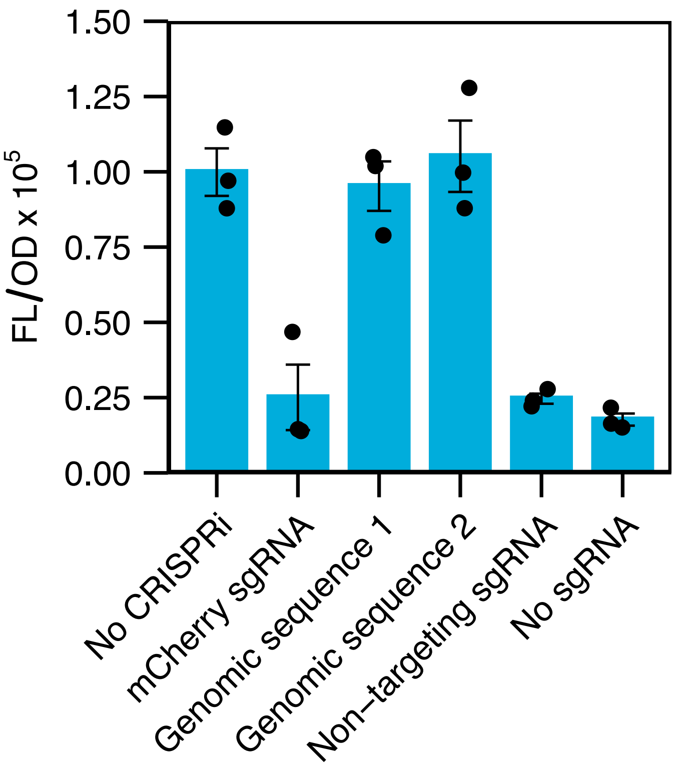

**Supplementary figure 2. CRISPRi results in decreased fluorescence in the absence of a sgRNA or in the presence of non-targeting sgRNA.** Fluorescence characterization of *S. venezuelae* cells containing a genomically-integrated mCherry gene conjugated with CRISPRi plasmids containing different sgRNAs or a no CRISPRi control plasmid. mCherry sgRNA binds to the mCherry gene and represses expression. Genomic sequence 1 and 2 are sgRNAs designed to non-coding sequences present in the *S. venezuelae* genome. Non-targeting sgRNA is designed to target a sequence absent in the *S. venezuelae* genome. No sgRNA is a CRISPRi plasmid without an sgRNA. Fluorescence characterization was performed by bulk fluorescence measurements (measured in units of fluorescence [FL]/optical density [OD] at 600 nm). Data represent mean values and errors bars represent standard deviation of at least 3 biological replicates.

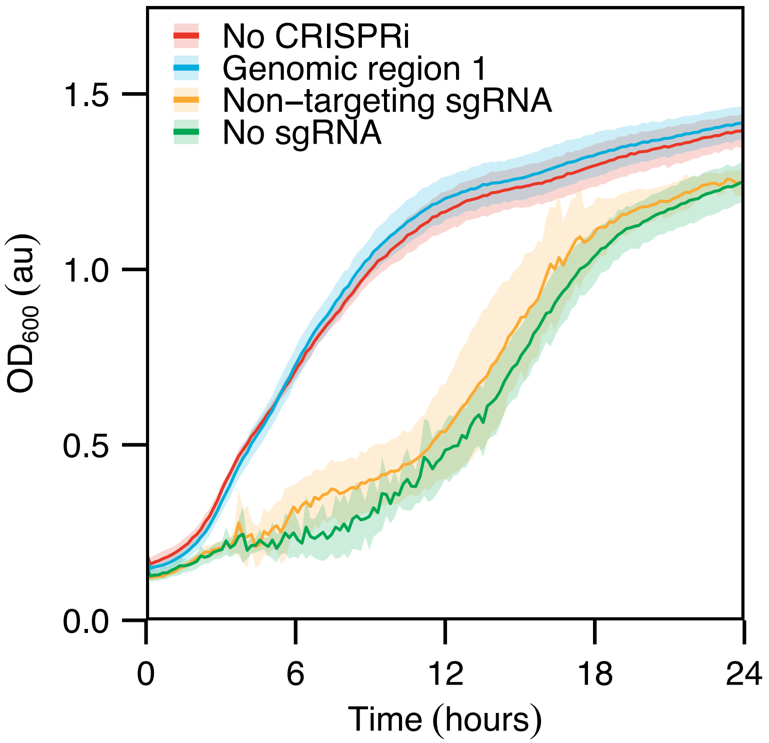

**Supplementary figure 3. CRISPRi results in inhibition of growth in the absence of a sgRNA or in the presence of non-targeting sgRNA.** Growth measurement of *S. venezuelae* conjugated with CRISPRi plasmids containing different sgRNAs or a no CRISPRi control plasmid. mCherry sgRNA binds to the mCherry gene and represses expression. Genomic region is a sgRNA designed to a non-coding sequence present in the *S. venezuelae* genome. Non-targeting sgRNA is designed to target a sequence absent in the *S. venezuelae* genome. No sgRNA is a CRISPRi plasmid without an sgRNA. Growth was evaluated by measuring optical density at 600 nm ($OD_{600}$) every 10 minutes at 30 °C with 90 rpm shaking. Lines represent mean values over time and shaded area depicts standard deviation of 4 biological replicates.

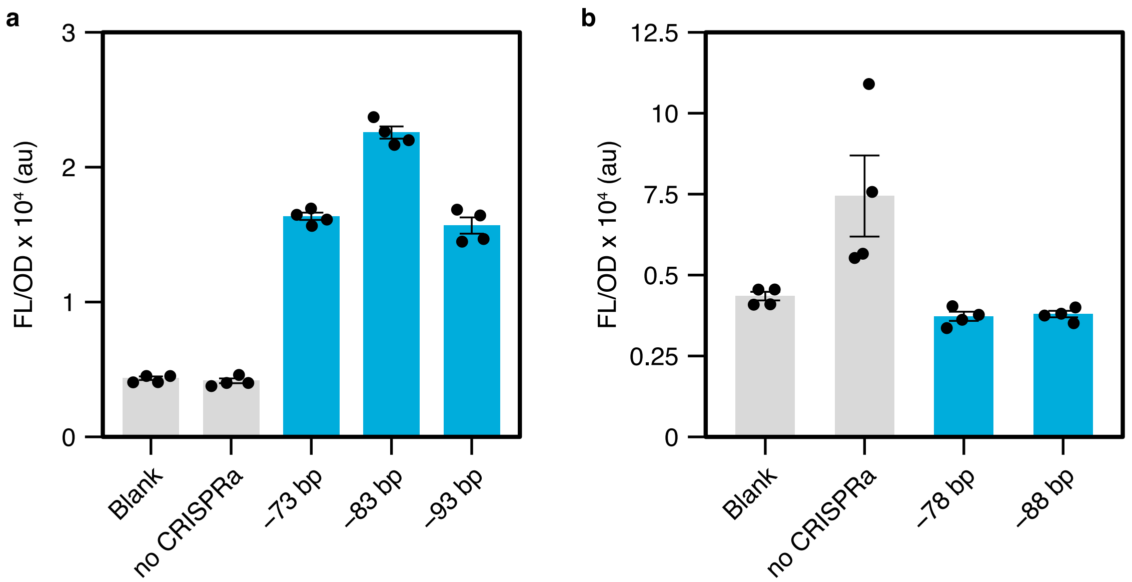

**Supplementary figure 4. Evaluating distance-dependent activation patterns of CRISPRa.** Fluorescence characterization of *S. venezuelae* cells containing a genomically-integrated mCherry gene conjugated with CRISPRa plasmids containing different sgRNAs or a no CRISPRa control plasmid. Blank cells are *S. venezuelae* cells lacking mCherry and transformed with the no CRISPRa plasmid that are used to determine autofluorescence. In (a) a reporter containing PAMs on the template strand at 73, 83, and 93 bp upstream of the reporter promoter's TSS is used. In (b) an additional 5 bp are added before the reporter promoter's TSS to create PAMs at 78 and 88 bp upstream. Fluorescence characterization was performed by bulk fluorescence measurements (measured in units of fluorescence [FL]/optical density [OD] at 600 nm). Data represent mean values and errors bars represent standard deviation of at least 4 biological replicates

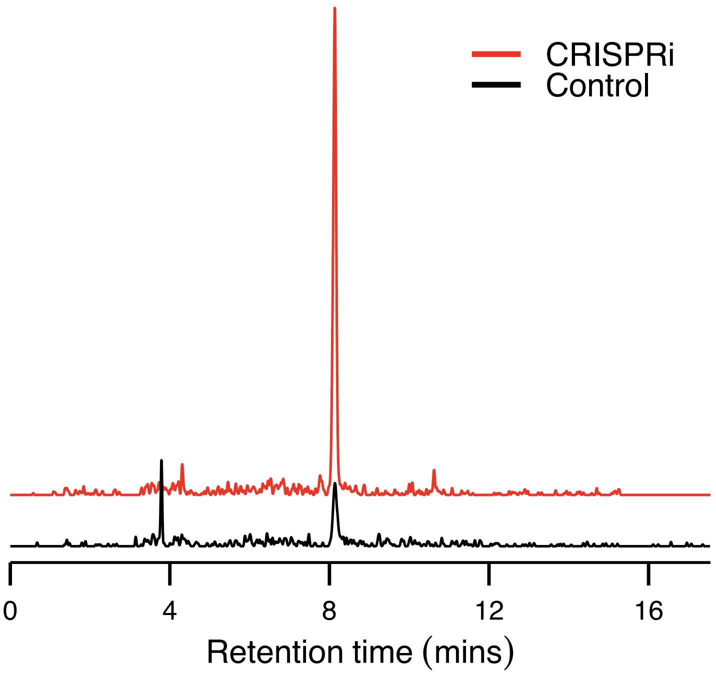

**Supplementary figure 5. Activating production of jadomycin B using CRISPRi.** LC-MS analysis of crude extracts of *S. venezuelae* cells conjugated with CRISPRi plasmids designed to repress the expression of jadR2. Cells were cultured, fermented, and extracted as described in methods. The crude extracts were then analyzed via liquid chromatography coupled to mass spectrometry (LC-MS). The reported data are extracted ion chromatogram at the corresponding *m/z* value of jdB (m/z = 550.2059, $[M+H]^+$). A jdB standard was run in parallel, and showed the same elution time as the CRISPRi sample. We note that the observed elution time of these replicates differs from the data shown in Figure 3c due to different instrumentation. For these samples, LC-MS analysis was carried out using a Shimadzu IT-ToF MS system interfaced to a Shimadzu LC-20 ADXR LC system through an Electro Sprat Ionization (ESI) source. Separations were carried out using an Ascentis Express C18 column (1.0 mm ID x 150 mm with 90A, 2.7u particles). The LC was operated at a flow rate of 0.175 mL/min with mobile phase A (0.1% Formic Acid in Water) and B (0.1% Formic Acid in Acetonitrile). The Analytical separation was carried out over 10 min; going from 10%B (t= 0 min) to 95% B(t= 10 min).

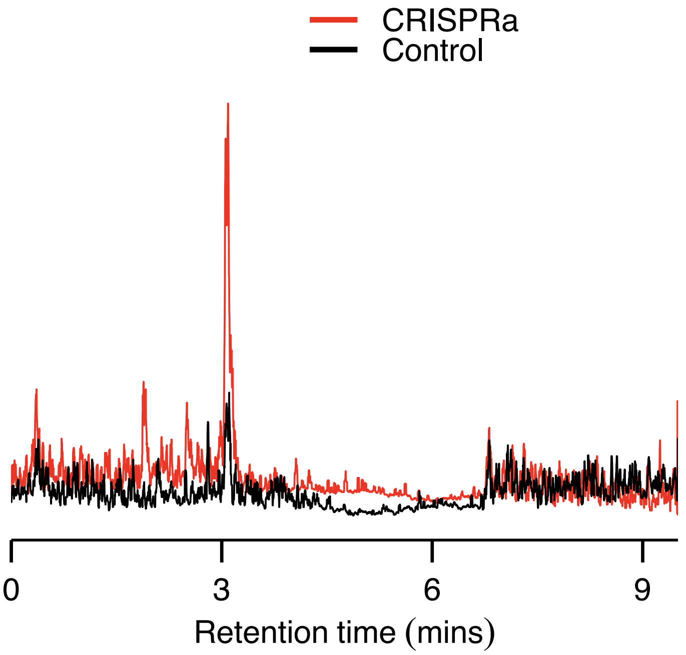

**Supplementary figure 6. Activating production of jadomycin B using CRISPRa.** LC-MS analysis of crude extracts of strains harboring jadJ-V-targeting CRISPRa plasmids. To induce the production of jdB, a sgRNA was designed to target a sequence downstream of a PAM site at 73 bp from the TSS of the jadJ-V operon within the jdB BGC. The sgRNA was cloned into a plasmid harboring the αNTD-based CRISPRa system. The plasmid was conjugated into wild-type *S. venezuelae*, and the resulting strains were cultured, fermented, and extracted. The crude extracts were then analyzed via liquid chromatography coupled to mass spectrometry (LC-MS), as described in the methods. The reported data are extracted ion chromatogram at the corresponding m/z value of jdB (m/z = 550.2059, $[M+H]^+$) for a second set of representative biological replicates (other than the ones shown in Figure 3d). A jdB standard was run in parallel, and showed the same elution time as the CRISPRa sample.
